# RAS–PI3K signaling promotes myofibroblastic CAF identity and restrains an immunomodulatory stromal program

**DOI:** 10.1101/2025.01.20.633776

**Authors:** Cristina Cuesta, Marta Alcón-Pérez, Rosa Ramírez-Cota, Sara Coadour, Jie Zheng, Nicole Procel, Dirk Fennema Galparsoro, Alejandro Rosell, Diana Loa-Meson, Belén Martínez-Castedo, Youssef Arafat, Héctor Sanz-Fraile, Vinothini Rajeeve, Diego Alonso-López, Robert E. Hynds, Charles Swanton, Jordi Alcaraz, Pedro Cutillas, Constantino C. Reyes-Aldasoro, Haiyun Wang, Esther Castellano

## Abstract

Cancer-associated fibroblasts display extensive functional plasticity, yet the signaling mechanisms that stabilize distinct CAF states remain poorly understood. Here, we identify stromal RAS–PI3K signaling as a key regulator of myofibroblastic CAF identity and matrix competence in lung cancer. Genetic disruption of the RAS–PI3K interaction in fibroblasts preserved upstream responsiveness to activating cues but impaired cytoskeletal remodeling, contractile execution and acquisition of a matrix-remodeling CAF program. As a result, RAS–PI3K-deficient CAFs generated extracellular matrices with defective collagen and fibronectin organization, reduced stiffness, altered microstructural properties and broad compositional remodeling of the matrisome, including changes in ECM glycoproteins and glycosylation-associated enzymes. These altered matrices failed to efficiently provide directional, adhesive and proliferative cues to tumor cells, attenuated EMT-associated programs and increased sensitivity to cisplatin plus pemetrexed. *In vivo*, fibroblast-specific RAS–PI3K disruption reduced tumor burden, decreased tumor cell proliferation, prolonged survival and improved chemotherapy response. Loss of RAS–PI3K signaling did not simply suppress CAF activation, but redirected CAFs toward a STAT3/NF-κB-linked immunomodulatory state associated with *IL6*, *CXCL12* and *PDGFRα* induction. Functionally, this altered stromal state promoted iNOS-associated macrophage features and was accompanied *in vivo* by reduced macrophage accumulation and increased CD8⁺ T cell infiltration. These findings establish RAS–PI3K signaling as a stromal state-control mechanism that couples myofibroblastic matrix competence to tumor-supportive microenvironmental function while restraining alternative immunomodulatory CAF programs.

## INTRODUCTION

Cancer progression is governed not only by tumor cell-intrinsic alterations but also by dynamic interactions between malignant cells and the surrounding tumor microenvironment. Among stromal components, cancer-associated fibroblasts (CAFs) have emerged as key regulators of tumor growth, invasion, immune modulation and therapeutic response^1,2^. Rather than acting as passive structural elements, CAFs integrate tumor-derived, mechanical and inflammatory cues to shape extracellular matrix (ECM) architecture, paracrine signaling and immune cell organization within tumors^3-6^.

A central feature of CAF biology is functional plasticity. Single-cell, spatial and multiomic studies have revealed that CAFs comprise heterogeneous yet interconvertible states with distinct functions across tumor types^3,4,6-8^. Among the best-characterized populations, myofibroblastic CAFs (myCAFs) are associated with contractility, ECM production and mechanical remodeling, whereas inflammatory or immunomodulatory CAF states produce cytokines and chemokines that influence immune cell recruitment and function^3,4^. Additional CAF phenotypes, including antigen-presenting and tissue-specific fibroblast states, further contribute to the complexity of the stromal compartment^7-10^. Importantly, these states are not fixed lineages but dynamic functional programs shaped by local cues within the tumor ecosystem^3,11-13^. However, the signaling mechanisms that stabilize CAFs in a matrix-remodeling state or allow transition toward alternative immunomodulatory programs remain incompletely understood.

The ECM is one of the major functional outputs of CAF activity. In tumors, CAF-derived matrices are not merely structural scaffolds, but instructive microenvironmental compartments that regulate cell adhesion^14,15^, migration^16-19^, proliferation^19,20^, survival^21,22^, mechanotransduction^23^ and immune cell localization^24-26^. MyCAFs contribute to these functions by generating actomyosin-dependent contractile forces, depositing structural ECM components such as collagens and fibronectin, and remodeling matrix architecture, stiffness and composition^5,27-30^. Recent work has further emphasized that ECM function depends not only on the abundance of individual matrix proteins, but also on their assembly, maturation, post-translational modification and organization into higher-order structures^31,32^. Thus, defining how CAFs acquire and maintain matrix competence is essential to understanding how stromal cells shape tumor behavior.

The functional impact of CAFs is highly context-dependent, as illustrated by therapeutic strategies aimed at targeting the tumor stroma. Approaches that broadly deplete fibroblasts or disrupt stromal programs have produced mixed, and in some cases detrimental, outcomes, in part because CAF populations can exert both tumor-promoting and tumor-restraining functions^1,33-35^. These findings have shifted the field toward a more selective view of stromal therapy, in which the goal is not to eliminate CAFs indiscriminately, but to understand and modulate specific tumor-supportive CAF functions. Identifying signaling pathways that control CAF state balance and matrix competence is therefore a critical step toward rational stromal reprogramming.

RAS signaling is a central driver of cancer progression, best known for its tumor cell-intrinsic oncogenic functions. However, increasing evidence indicates that RAS pathway activity also shapes the tumor microenvironment by regulating paracrine signaling, immune evasion, angiogenesis and stromal remodeling^36-39^. In several tumor models, oncogenic KRAS has been shown to influence fibroblast activation and ECM remodeling through tumor cell-derived signals^39-42^. In contrast, the role of wild-type RAS signaling within stromal cells remains comparatively less explored. This distinction is important because stromal cells are genetically non-transformed but continuously exposed to tumor-derived cues that require interpretation through intracellular signaling networks.

PI3K is a major RAS effector^43^ with established roles in cytoskeletal organization, adhesion, contractility and cellular responses to mechanical cues^44-49^. Previous work showed that disrupting the interaction between RAS and the p110α catalytic subunit of PI3K abolishes lung tumor initiation and reduces lung tumor maintenance, macrophage recruitment and tumor-induced angiogenesis^50-52^, suggesting that RAS–PI3K signaling contributes to tumor progression beyond tumor cell-autonomous pathways. However, whether RAS–PI3K signaling in CAFs controls stromal functional identity, ECM assembly and tumor microenvironment organization remains unknown.

Here, we investigate how selective disruption of RAS–PI3K signaling in fibroblasts affects CAF state, matrix competence and tumor progression in KRAS-driven lung cancer. We show that RAS–PI3K signaling is required for CAFs to execute a myofibroblastic program, coupling cytoskeletal remodeling to ECM production, maturation and organization. Loss of this pathway does not simply suppress CAF activation, but redirects CAFs toward an alternative STAT3/IL6-linked immunomodulatory state. Functionally, RAS–PI3K-deficient CAFs generate structurally, mechanically and compositionally altered matrices that fail to efficiently support tumor cell organization, growth and chemoprotection, while fibroblast-specific RAS–PI3K disruption remodels the immune contexture of lung tumors in vivo. These findings identify stromal RAS–PI3K signaling as a regulator of CAF state balance and reveal matrix competence as a key mechanism through which CAFs shape the physical, immune and therapeutic landscape of the tumor microenvironment.

## RESULTS

### RAS–PI3K signaling is required to sustain CAF activation and ECM deposition *in vivo*

RAS–PI3K signaling regulates different aspects of fibroblast biology and has been implicated in lung tumor progression^48,50,52,53^. We therefore hypothesized that RAS–PI3K activity within CAFs is required to sustain their activated functional phenotype and that disruption of this signaling axis may contribute to the impaired tumor growth previously observed upon RAS–PI3K uncoupling.

To test this hypothesis *in vivo*, we used the *Pik3ca*^RBD^ mouse model, in which the interaction between p110α and RAS is selectively disrupted by two-point mutations in the endogenous *Pik3ca* gene^50,52^. Wild-type (*Pik3ca^WT^*) and *Pik3ca^RBD^* mice were bred with mice carrying floxed *Pik3ca* alleles and *Cre-ERT2* recombinase alleles targeted to the *Rosa26* locus (*Pik3ca^RBD/flox^*) and further crossed with Kras^LA2^ mice^54^ to generate lung tumors in which RAS–PI3K signaling can be specifically disrupted following tamoxifen administration^50^. After 16 weeks of tamoxifen treatment, lungs were harvested and analyzed for CAF activation markers. Histological analysis of lung tumors revealed a marked reduction in CAF activation upon RAS–PI3K disruption. α-SMA immunostaining showed a substantial decrease in α-SMA–positive stromal cells in *Pik3ca^RBD/-^*tumors compared to *Pik3ca^WT/-^* controls (Fig. 1A). Consistent with this, Sirius Red staining demonstrated a pronounced reduction in fibrillar collagen deposition in *Pik3ca^RBD/-^* tumors, indicating impaired ECM accumulation in the absence of RAS–PI3K signaling (Fig. 1A). These data suggest that disruption of RAS–PI3K signaling may alter the functional state of the stromal compartment.

**Figure 1.**
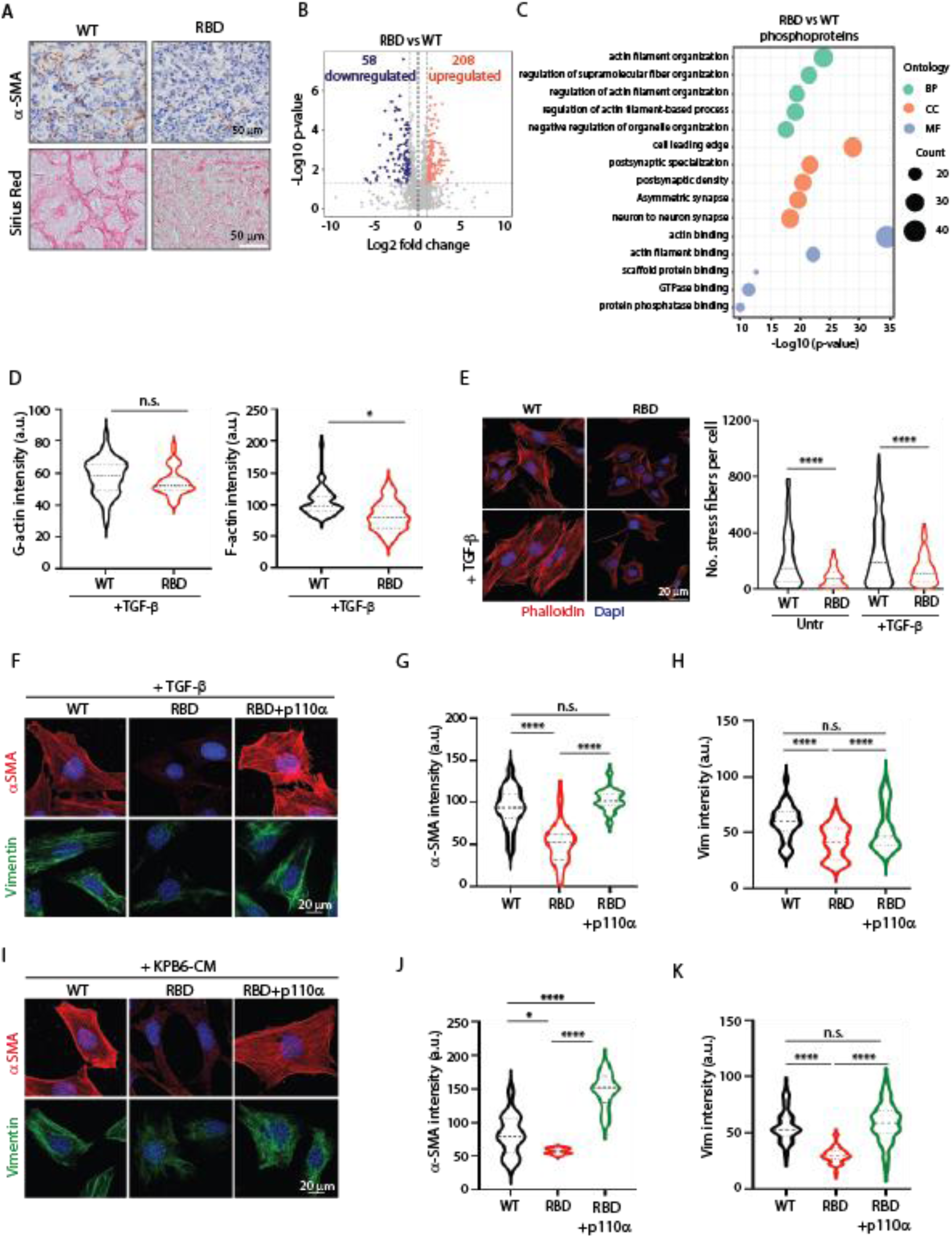
RAS–PI3K signaling is required for cytoskeletal remodeling and myCAF activation. (A) Representative images of α-SMA immunohistochemistry and Sirius Red staining in lung tumors from *Pik3ca*^WT/−^ and *Pik3ca*^RBD/−^ mice, showing reduced stromal α-SMA expression and fibrillar collagen deposition upon disruption of the RAS–PI3K interaction. Scale bars, 50 μm. (B) Volcano plot showing differentially phosphorylated peptides identified by phosphoproteomic analysis of *Pik3ca*^RBD/−^ *versus Pik3ca*^WT/−^ lung tumors. Significantly downregulated and upregulated phosphopeptides are shown in blue and red, respectively. (C) Gene Ontology enrichment analysis of significantly altered phosphopeptides in *Pik3ca*^RBD/−^ *versus Pik3ca*^WT/−^ tumors, highlighting enrichment of processes related to actin filament organization, cytoskeletal regulation, cell polarity and adhesion. Dot size indicates the number of proteins associated with each term; color denotes ontology category. (D) IF-based quantification of G-actin and F-actin levels in WT and RBD fibroblasts following TGF-β stimulation, showing impaired actin polymerization in RAS–PI3K-deficient fibroblasts. (E) Representative phalloidin staining and quantification of stress fiber formation in WT and RBD fibroblasts under basal conditions or after TGF-β stimulation. Phalloidin, red; DAPI, blue. Scale bar, 20 μm. (F) Representative immunofluorescence images of α-SMA and vimentin in WT, RBD and RBD fibroblasts re-expressing functional p110α following TGF-β stimulation. α-SMA, red; vimentin, green; DAPI, blue. Scale bar, 20 μm. (G–H) Quantification of α-SMA (G) and vimentin (H) fluorescence intensity following TGF-β stimulation. (I) Representative immunofluorescence images of α-SMA and vimentin in WT, RBD and RBD+p110α fibroblasts stimulated with conditioned medium from KRAS-mutant KPB6 lung cancer cells. α-SMA, red; vimentin, green; DAPI, blue. Scale bar, 20 μm. (J–K) Quantification of α-SMA (J) and vimentin (K) fluorescence intensity in the conditions shown in (I). Data are shown as violin plots with median and quartiles. Statistical significance was determined using Student’s t-test unless otherwise indicated. p < 0.05; ***p < 0.0001; n.s., not significant.

To gain insight into the signaling landscape associated with RAS–PI3K disruption *in vivo*, we performed phosphoproteomic profiling of tumors from *Pik3ca^RBD/-^* and *Pik3ca^WT/-^* mice. To assess whether these signaling changes could be recapitulated pharmacologically, we additionally analyzed tumors from wild-type mice treated with the p110α-specific inhibitor alpelisib. We identified 266 differentially phosphorylated peptides in *Pik3ca^RBD/-^* tumors compared to controls, with 208 upregulated and 58 downregulated by at least twofold (Fig. 1B and Supplementary Table S1). In alpelisib-treated mice, we identified a total of 241 differentially phosphorylated peptides compared to controls (68 downregulated and 173 upregulated) (Supplementary Fig. S1A). Notably, a substantial proportion of phosphoproteomic changes overlapped between *Pik3ca^RBD/-^* tumors and alpelisib-treated tumors, with 163 proteins showing concordant regulation across both conditions (Supplementary Fig. S1B), supporting the robustness of RAS–PI3K-dependent signaling alterations *in vivo*.

As this approach captures integrated signaling alterations across multiple cellular compartments within the tumor, pathway-level analysis was used to interpret these changes. Ontology enrichment analysis highlighted alterations in pathways governing actin filament organization, cellular polarity, and cell–cell adhesion, both in *Pik3ca^RBD/-^* tumors (Fig. 1C) and in alpelisib treated mice (Supplementary Fig. S1C). These processes are particularly relevant for stromal cell populations that rely on dynamic cytoskeletal remodeling to acquire and maintain an activated state^5,28^.

Based on this convergence on cytoskeletal pathways, we next asked whether RAS–PI3K signaling is required for the implementation of actin remodeling programs that are central to CAF activation and function. To directly test this possibility in a CAF-relevant context, we isolated embryonic fibroblasts from *Pik3ca*^WT^ and *Pik3ca*^RBD^ mice^52^ and examined their ability to reorganize the actin cytoskeleton upon stimulation with transforming growth factor β (TGF-β), a key profibrotic signal that drives CAF activation *in vitro*. Biochemical fractionation of globular (G-) and filamentous (F-) actin revealed a marked reduction in F-actin levels in RAS–PI3K-deficient fibroblasts following TGF-β stimulation, while G-actin levels remained unchanged (Fig. 1D).

To assess whether this defect translated into altered cytoskeletal organization at the cellular level, we next analyzed stress fiber formation and overall actin architecture in activated fibroblasts. Phalloidin staining revealed a pronounced reduction in stress fiber formation in RAS–PI3K-deficient fibroblasts compared with wild-type controls upon TGF-β stimulation (Fig. 1E). Quantitative analysis confirmed a significant decrease in stress fiber abundance and organization, consistent with defective actin remodeling downstream of RAS–PI3K disruption. In line with these cytoskeletal defects, RAS–PI3K-deficient fibroblasts also exhibited altered cell morphology, as reflected by increased cellular circularity following TGF-β stimulation (Supplementary Fig. S1D).

Given the defects in actin polymerization, stress fiber formation, and cell morphology observed upon RAS–PI3K disruption, we next asked whether these cytoskeletal alterations translated into impaired execution of an active fibroblastic CAF program. To this end, we examined the induction of α-SMA and vimentin in fibroblasts stimulated with either TGF-β or conditioned medium derived from KRAS-mutant lung cancer cells (KPB6), another protumorigenic cue known to promote CAF activation. Immunofluorescence analysis revealed a significant decrease of α-SMA and vimentin in RAS–PI3K-deficient fibroblasts (Fig. 1F–K). Importantly, re-expression of functional p110α in RBD fibroblasts restored α-SMA and vimentin expression following stimulation with either TGF-β or KPB6-CM, confirming that these defects are specifically linked to disruption of the RAS–PI3K axis. Consistent with the immunofluorescence data, immunoblot analysis confirmed a marked reduction in α-SMA and vimentin protein levels in RAS–PI3K-deficient fibroblasts following stimulation with TGF-β or conditioned medium (Supplementary Fig. S1E).

Canonical TGF-β signaling remained largely intact in RAS–PI3K-deficient fibroblasts, as evidenced by comparable SMAD2/3 phosphorylation dynamics (Supplementary Fig. S1F) and similar levels of surface TGF-β receptor expression (Supplementary Fig. S1G), indicating preserved signal perception. In contrast, AKT phosphorylation was selectively reduced, while ERK activation was unaffected (Supplementary Fig. S1H), consistent with effective disruption of the RAS–PI3K axis.

Together, these data demonstrate that RAS–PI3K signaling is essential for the functional deployment of cytoskeletal remodeling programs required for CAF activation in response to both profibrotic and tumor-derived cues.

### RAS–PI3K signaling shapes CAF transcriptional identity

These observations indicate that RAS–PI3K signaling is required to sustain key functional features of activated CAFs. However, it remains unclear whether loss of this pathway simply attenuates CAF activation or instead reprograms CAF transcriptional identity.

To address this, we performed transcriptomic profiling of WT and RAS–PI3K-deficient CAFs stimulated with either TGF-β or conditioned medium from KRAS-mutant lung cancer cells (KPB6). Unsupervised multidimensional scaling (MDS) analysis revealed that the primary axis of variation segregated samples according to RAS–PI3K status, whereas the activating stimulus contributed to a secondary axis of variation, suggesting that loss of RAS–PI3K signaling establishes a distinct transcriptional state that shapes CAF responses to protumorigenic cues (Fig. 2A).

**Figure 2.**
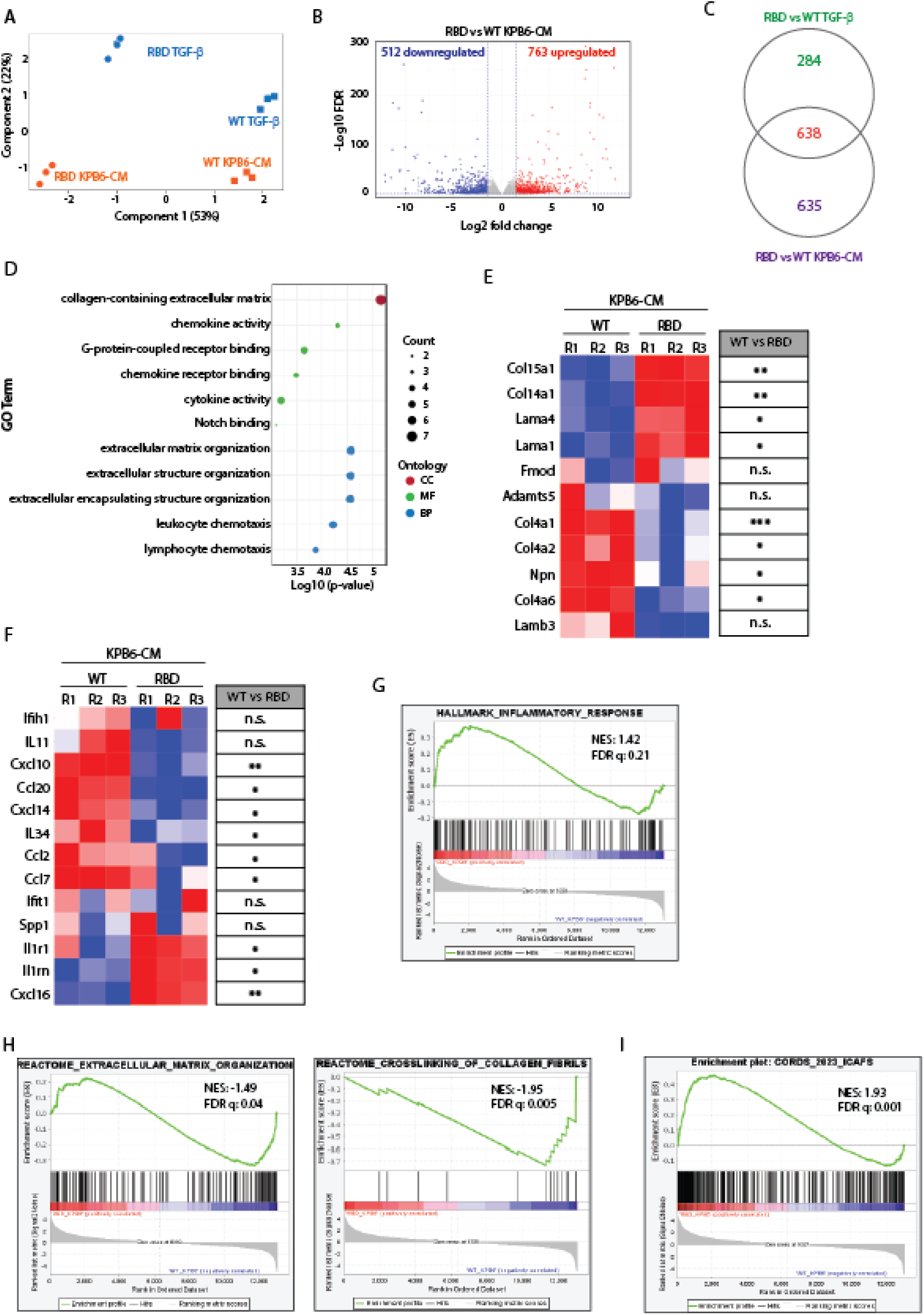
RAS–PI3K signaling controls the balance between matrix-remodeling and immunomodulatory CAF programs. (A) Multidimensional scaling analysis of RNA-seq profiles from WT and RAS–PI3K-deficient RBD CAFs stimulated with TGF-β or conditioned medium from KRAS-mutant KPB6 lung cancer cells. Samples segregate primarily according to RAS–PI3K status, with activating stimulus contributing to a secondary axis of variation. (B) Volcano plot showing differentially expressed genes in RBD *versus* WT CAFs following stimulation with KPB6-CM. Significantly downregulated and upregulated genes are shown in blue and red, respectively. (C) Venn diagram showing overlap between differentially expressed genes identified in RBD *versus* WT CAFs following TGF-β or KPB6-CM stimulation. (D) Gene Ontology enrichment analysis of shared differentially expressed genes, highlighting extracellular matrix–associated and immune-related biological processes. Dot size indicates the number of genes associated with each term; color denotes ontology category. (E) Heatmap showing quantitative PCR validation of selected extracellular matrix–associated genes in WT and RBD CAFs following KPB6-CM stimulation. Statistical significance for WT versus RBD comparisons is indicated on the right. For heatmaps, each column represents an independent biological replicate. (F) Heatmap showing quantitative PCR validation of selected immune-related and cytokine/chemokine-associated genes in WT and RBD CAFs following KPB6-CM stimulation. Statistical significance for WT versus RBD comparisons is indicated on the right. (G) GSEA showing enrichment of the HALLMARK_INFLAMMATORY_RESPONSE signature in RBD CAFs compared with WT CAFs following KPB6-CM stimulation. (H) GSEA showing enrichment of extracellular matrix organization and collagen fibril crosslinking signatures in WT CAFs compared with RBD CAFs following KPB6-CM stimulation. (I) Gene Set Enrichment Analysis showing enrichment of a published inflammatory CAF signature in RBD CAFs compared with WT CAFs under tumor-conditioned medium stimulation. Statistical significance was determined using Student’s t-test unless otherwise indicated. *p < 0.05; **p < 0.01; ***p < 0.001; n.s., not significant. GSEA plots show normalized enrichment score (NES) and false discovery rate (FDR q value).

Differential expression analysis revealed extensive transcriptional changes in RBD CAFs relative to WT controls in response to both stimuli (Fig. 2B and supplementary Fig. S2A). Upon TGF-β stimulation, 440 genes were upregulated and 482 downregulated in RBD versus WT CAFs (Supplementary Fig. S2A and Supplementary table S3), whereas KPB6-CM elicited a significantly broader response (763 upregulated and 512 downregulated genes) (Fig. 2B and Supplementary table S4), consistent with the more complex nature of tumor-derived conditioned medium. Heatmap visualization further supported a robust separation between WT and RBD transcriptional programs under each condition (Supplementary Fig. S2B).

To assess whether these transcriptional changes reflected stimuli-specific effects or a common program associated with RAS–PI3K loss, we compared differentially expressed genes across both conditions. This analysis revealed a substantial overlap between the transcriptional responses induced by TGF-β and KPB6-CM (638 shared genes) (Fig. 2C). Given that tumor-derived conditioned medium contains TGF-β among other protumorigenic factors, this overlap likely reflects the engagement of shared signaling inputs that are interpreted through a RAS–PI3K-dependent transcriptional framework. Notably, shared TGF-β and KPB6-CM DEGs displayed concordant directionality of regulation across both conditions (Supplementary Fig. S2C-D), indicating that loss of RAS–PI3K signaling imposes a common transcriptional bias on CAFs exposed to distinct, but partially overlapping, protumorigenic cues.

Gene Ontology enrichment analysis of shared DEGs identified two dominant functional modules: extracellular matrix–related processes, including collagen-containing ECM and structural organization, and immune-related categories associated with cytokine activity and leukocyte chemotaxis (Fig. 2D).

Consistent with the transcriptomic analysis, quantitative PCR validation of selected genes in both modules supported the presence of coordinated changes in both extracellular matrix–related and immunomodulatory programs in RAS–PI3K-deficient CAFs following stimulation with KPB6-CM (Fig. 2E–F). ECM-associated genes identified in the RNA-seq dataset, including collagens and basement membrane components such as *Col15a1*, *Col14a1*, *Lama1* and *Lama4*, showed an overall reduction in RAS–PI3K-deficient CAFs, with most reaching statistical significance across three independent biological replicates (Fig. 2E). Only *Adamts5*, *Fmod* and *Lamb3* did not reach statistical significance. In parallel, validation of immune-related genes revealed a selective rewiring of the cytokine and chemokine response to tumor-conditioned medium. Several tumor-induced inflammatory mediators, including *Cxcl10*, *Ccl20*, *Cxcl14*, *Il34*, *Ccl2* and *Ccl7*, were significantly reduced in RAS–PI3K-deficient CAFs, whereas *Il1r1*, *Il1rn* and *Cxcl16* were increased (Fig. 2F). Thus, loss of RAS–PI3K signaling does not induce a uniform inflammatory response, but instead redirects CAFs toward a distinct immunomodulatory transcriptional state.

To further contextualize these transcriptional changes within established CAF phenotypic frameworks, we performed Gene Set Enrichment Analysis (GSEA). RAS–PI3K-deficient CAFs exposed to KPB6-CM showed significant enrichment of the HALLMARK_INFLAMMATORY_RESPONSE signature (Fig. 2G), whereas WT CAFs were enriched for extracellular matrix organization and collagen fibril crosslinking signatures (Fig. 2H). These findings indicate that RAS–PI3K disruption shifts CAFs away from matrix-remodeling transcriptional programs and toward inflammatory/immunomodulatory responses.

We next interrogated recently defined CAF subtype signatures^7^. Under tumor-derived conditioned medium stimulation, RAS–PI3K-deficient CAFs showed significant enrichment of a published inflammatory CAF signature (Fig. 2I), further supporting a shift toward an iCAF-like transcriptional state. Together, these analyses indicate that loss of RAS–PI3K signaling does not simply attenuate CAF activation, but changes the balance between matrix-remodeling and immunomodulatory CAF programs.

### CAF-specific RAS–PI3K signaling sustains tumor growth and stromal organization *in vivo*

Given the transcriptional rewiring induced by RAS–PI3K loss in CAFs, we next asked whether these alterations translated into functional consequences for tumor progression in vivo. To address this, we generated a fibroblast-specific derivative of the *Pik3ca*^RBD/flox^ model in which disruption of the RAS–PI3K interaction was selectively induced in fibroblasts by Cre recombinase expression under the control of the *Col1a2* promoter (*Pik3ca*^fRBD/flox^) (Supplementary Fig. S3A). Tumor growth was evaluated using an experimental lung colonization model based on intravenous tail-vein injection of KRAS-mutant KPB6 cells, which preferentially seed and expand within the lung parenchyma.

Selective disruption of RAS–PI3K signaling in CAFs resulted in a significant reduction in tumor burden compared to control mice (Fig. 3A), indicating that RAS–PI3K activity in fibroblasts contributes to tumor progression in the context of aggressive tumor growth. To define the cellular basis of this effect, we examined proliferative activity within tumor lesions. Quantification of phospho-histone H3–positive nuclei revealed a significant decrease in mitotic figures in tumors developing in mice harboring RAS–PI3K-deficient CAFs (Fig. 3B). Consistently, immunohistochemical analysis showed decreased pERK staining within tumor regions, indicating reduced MAPK pathway activation (Fig. 3C), in line with impaired tumor cell proliferation.

**Figure 3.**
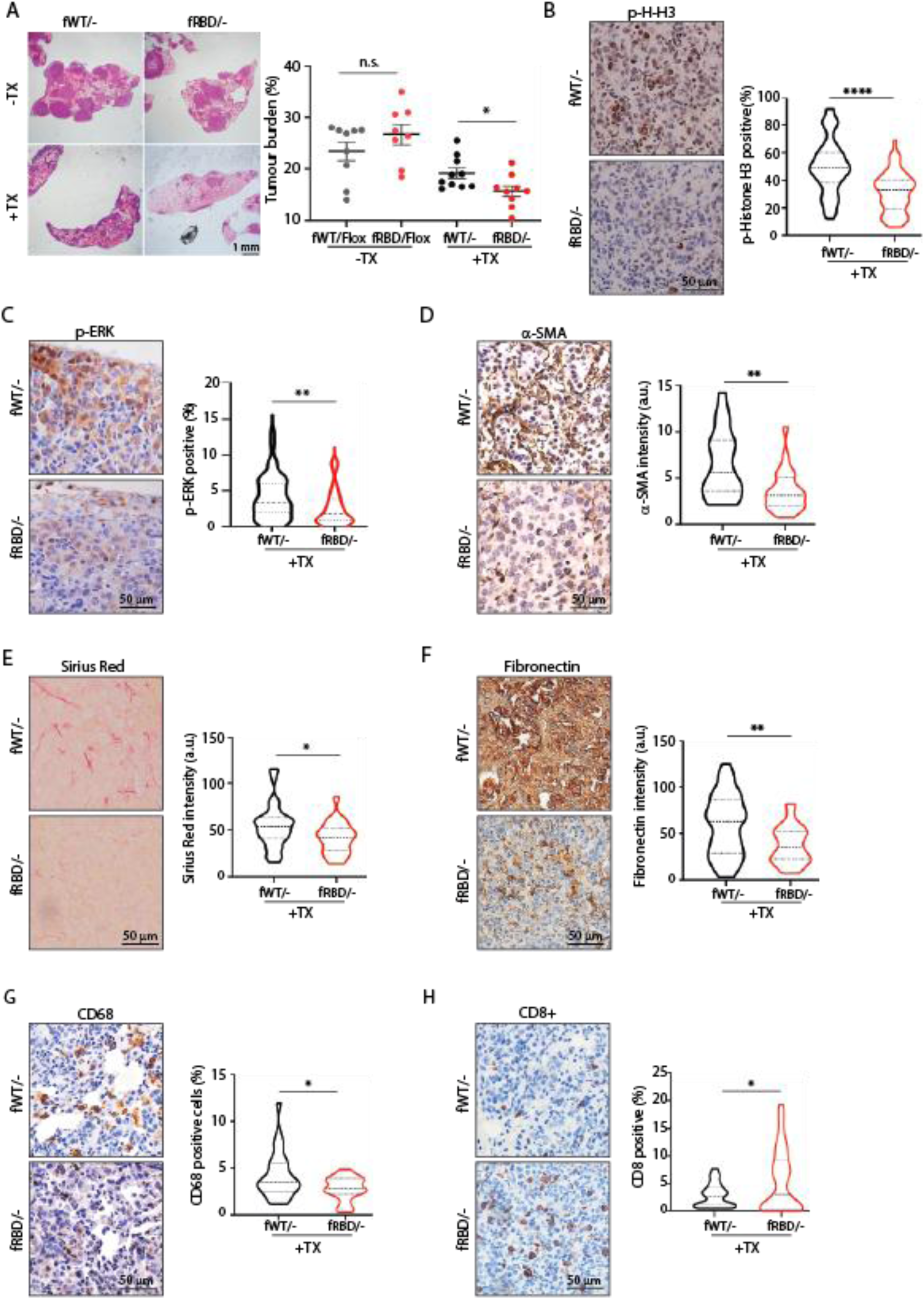
CAF-specific disruption of RAS–PI3K signaling restricts lung tumor growth and remodels the tumor microenvironment. (A) Representative H&E-stained lung sections and quantification of tumor burden in *Pik3ca*^fWT/−^ and *Pik3ca*^fRBD/-^ mice following intravenous injection of KRAS-mutant KPB6 lung cancer cells. Mice were analyzed in the absence or presence of tamoxifen-induced recombination (+TX). Tumor burden is expressed as tumor area relative to total lung area. Each dot represents an individual mouse. Scale bar, 1 mm. (B) Representative phospho-histone H3 immunohistochemistry and quantification of p-Histone H3–positive nuclei in tumors from tamoxifen-treated *Pik3ca*^fWT/−^ and *Pik3ca*^fRBD/-^ mice, showing reduced tumor cell proliferation upon CAF-specific RAS–PI3K disruption. Scale bar, 50 μm. (C) Representative pERK immunohistochemistry and quantification of pERK-positive tumor area in tumors from tamoxifen-treated *Pik3ca*^fWT/−^ and *Pik3ca*^fRBD/-^ mice. Scale bar, 50 μm. (D) Representative α-SMA immunohistochemistry and quantification of α-SMA staining in tumors from tamoxifen-treated *Pik3ca*^fWT/−^ and *Pik3ca*^fRBD/-^ mice, indicating reduced myofibroblastic CAF activation. Scale bar, 50 μm. (E) Representative Sirius Red staining and quantification of fibrillar collagen deposition in tumors from tamoxifen-treated *Pik3ca*^fWT/−^ and *Pik3ca*^fRBD/-^ mice. Scale bar, 50 μm. (F) Representative fibronectin immunohistochemistry and quantification of fibronectin deposition in tumors from tamoxifen-treated *Pik3ca*^fWT/−^ and *Pik3ca*^fRBD/-^ mice. Scale bar, 50 μm. (G) Representative CD68 immunohistochemistry and quantification of CD68-positive macrophages in tumors from tamoxifen-treated *Pik3ca*^fWT/−^ and *Pik3ca*^fRBD/-^ mice. Scale bar, 50 μm. (H) Representative CD8 immunohistochemistry and quantification of CD8-positive T cells in tumors from tamoxifen-treated *Pik3ca*^fWT/−^ and *Pik3ca*^fRBD/-^ mice. Scale bar, 50 μm. Data are shown as mean ± SEM in (A) and as violin plots with median and quartiles in (B–H). Statistical significance was determined using Student’s t-test unless otherwise indicated. p < 0.05; *p < 0.01; ***p < 0.0001; n.s., not significant.

Histological analysis further revealed that reduced tumor burden was accompanied by marked alterations in stromal organization. Tumors arising from the *Pik3ca*^fRBD/-^ mice exhibited a significant decrease in α-SMA–positive CAFs, together with reduced deposition of fibronectin and fibrillar collagen within tumor lesions compared to controls (Fig. 3D–F), consistent with the impaired matrix-remodeling programs identified in the RNAseq analysis.

Given the transcriptional enrichment of inflammatory and immune-related pathways in RAS–PI3K-deficient CAFs, we next examined whether stromal RAS–PI3K disruption was associated with changes in tumor immune composition. Tumors from *Pik3ca*^fRBD/−^ mice displayed reduced macrophage infiltration together with a significant increase in CD8⁺ T cell accumulation compared with controls (Fig. 3G–H), indicating selective remodeling of the tumor immune microenvironment. In contrast, CD4⁺ T cell levels were not significantly altered (Supplementary Fig. S3B), suggesting that CAF-specific RAS–PI3K disruption does not produce a uniform change in lymphocyte infiltration.

To validate these findings in an autochthonous setting, we next used a genetic lung cancer model. *Pik3ca*^fRBD/flox^ mice were crossed with KRAS^LA2^ animals to induce spontaneous lung tumor formation, and recombination was triggered by tamoxifen administration. Lungs were harvested and analyzed 16 weeks after tamoxifen treatment. Consistent with the lung colonization model, fibroblast-specific disruption of RAS–PI3K signaling resulted in a significant reduction in the number of tumors on lung surface (Supplementary Fig. S3C) and in overall tumor burden (Supplementary Fig. S3D). Histological and immunohistochemical analyses further revealed a marked decrease in α-SMA–positive stromal cells, reduced collagen deposition, and diminished fibronectin accumulation in tumors from *Pik3ca*^fRBD/-^ mice compared to *Pik3ca*^fWT/-^ controls (Supplementary Fig. S3E).

Together, these findings indicate that CAF-specific RAS–PI3K signaling supports lung tumor progression and stromal organization *in vivo*. Its disruption reduces tumor burden, decreases tumor cell proliferative signaling, impairs deposition of major ECM components and is associated with selective remodeling of immune cell composition. These alterations prompted us to investigate whether RAS–PI3K-deficient fibroblasts have an intrinsic defect in the production and organization of tumor-supportive extracellular matrices.

### RAS–PI3K signaling is required for CAF contractility and matrix assembly competence

A defining functional property of myCAFs is their ability to generate contractile forces that compact and remodel collagen-rich extracellular matrices^55^. Given the cytoskeletal defects observed upon RAS–PI3K disruption (Fig. 1), we examined whether RAS–PI3K-deficient fibroblasts retain the capacity to exert contractile tension in three-dimensional collagen gels. While WT fibroblasts efficiently contracted collagen matrices upon TGF-β stimulation, RAS–PI3K-deficient fibroblasts presented a marked impairment in gel contraction ability (Fig. 4A-B). Reintroduction of functional p110α into RBD cells restored contractile capacity to WT levels, confirming that this defect is attributable to disruption of the RAS–PI3K axis. Pharmacological inhibition of PI3Kα with alpelisib or with a RAS–PI3K Breaker (Supplementary Fig. S4A), a small molecule that uncouples the RAS–PI3K interaction^56^, similarly impaired gel contraction in WT fibroblasts, phenocopying the genetic RAS–PI3K-deficient state (Fig. 4A-B).

**Figure 4.**
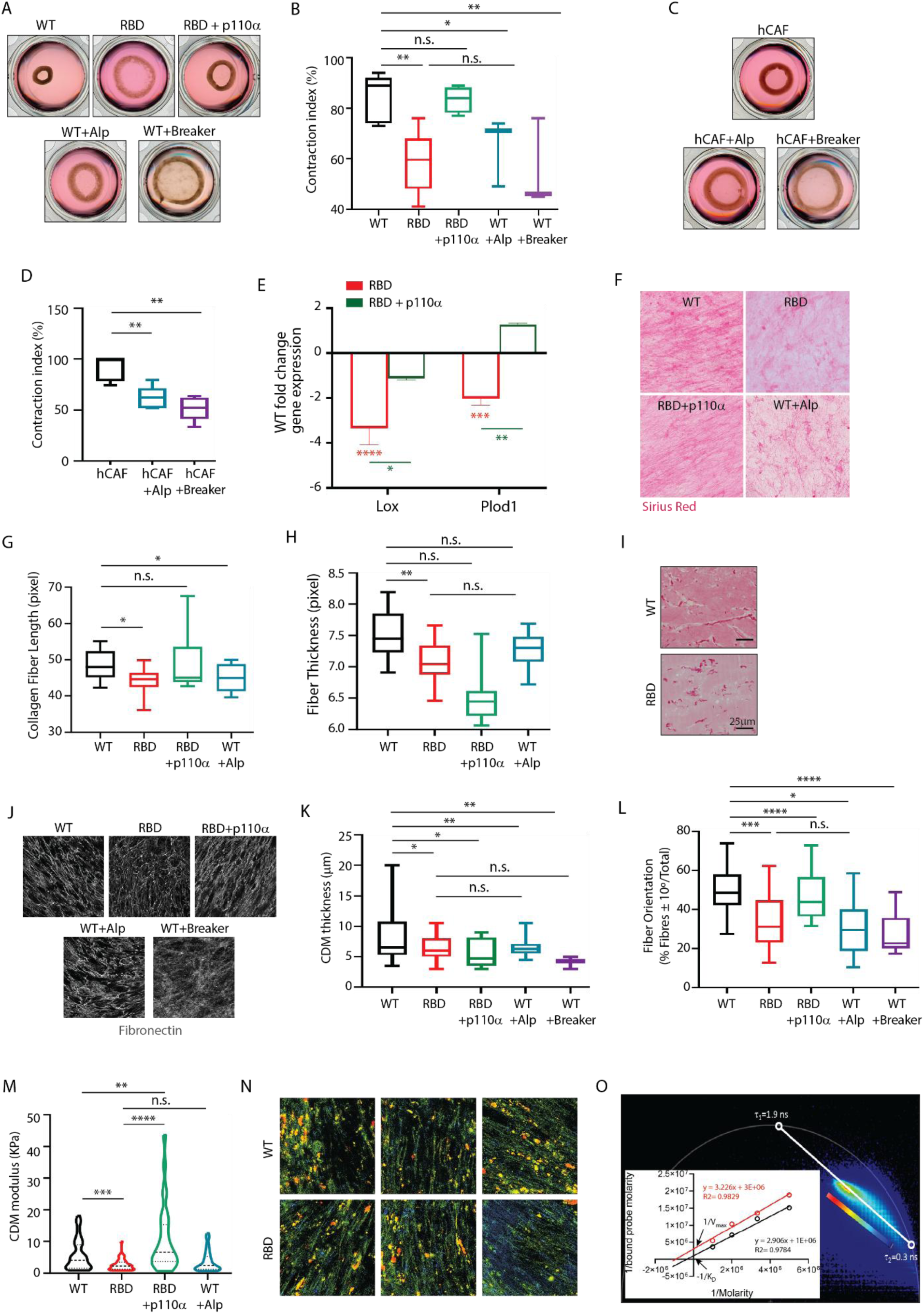
RAS–PI3K signaling is required for CAF contractility and extracellular matrix assembly competence. (A) Representative images of three-dimensional collagen gels containing WT, RBD or RBD+p110α fibroblasts, or WT fibroblasts treated with alpelisib or the RAS–PI3K interaction Breaker. Gels were imaged after contraction. (B) Quantification of collagen gel contraction index for the conditions shown in (A), showing impaired collagen contraction upon genetic or pharmacological disruption of RAS–PI3K signaling and rescue following re-expression of functional p110α. (C) Representative images of collagen gels containing human lung cancer-associated fibroblasts (hCAFs) treated with vehicle, alpelisib or Breaker. (D) Quantification of collagen gel contraction index in hCAFs, showing reduced contractile capacity upon pharmacological inhibition of PI3Kα or disruption of RAS–PI3K coupling. (E) Quantitative PCR analysis of Lox and Plod1 expression in RBD versus WT fibroblasts, and in RBD+p110α versus RBD fibroblasts, following stimulation with KPB6-CM. Data are shown as fold change relative to WT controls. (F) Representative Sirius Red staining of CAF-derived matrices generated by WT, RBD, RBD+p110α or alpelisib-treated WT fibroblasts. (G–H) Quantification of collagen fiber length (G) and fiber thickness (H) in CAF-derived matrices shown in (F). (I) Representative Sirius Red staining of collagen plugs containing TGF-β-activated WT or RBD fibroblasts following implantation in syngeneic mice, showing impaired collagen organization by RAS–PI3K-deficient fibroblasts *in vivo*. Scale bar, 25 μm. (J) Representative fibronectin immunofluorescence images of CAF-derived matrices generated by WT, RBD or RBD+p110α fibroblasts, or WT fibroblasts treated with alpelisib or Breaker. (K–L) Quantification of matrix thickness (K) and fibronectin fiber orientation (L) in the conditions shown in (J). (M) Atomic force microscopy analysis of the elastic modulus of CAF-derived matrices generated by WT, RBD or RBD+p110α fibroblasts, or WT fibroblasts treated with alpelisib, showing reduced matrix stiffness upon genetic or pharmacological disruption of RAS–PI3K signaling.. (N) Representative fluorescence lifetime images of WT and RBD CAF-derived matrices stained with increasing concentrations of the molecular rotor DCVJ. Shorter fluorescence lifetimes indicate altered molecular restriction and microstructural organization within RBD-derived matrices. (O) Phasor-based FLIM analysis of DCVJ fluorescence lifetime distributions in WT and RBD CAF-derived matrices. The two lifetime components correspond to matrix-associated/restricted and freely diffusing DCVJ populations. Inset, phasor-derived binding analysis represented as a Lineweaver–Burk plot. Data are shown as box-and-whisker plots, violin plots or bar graphs as indicated. Statistical significance was determined using Student’s t-test unless otherwise indicated. *p < 0.05; **p < 0.01; ***p < 0.001; ****p < 0.0001; n.s., not significant.

Consistent with these findings, both alpelisib and the Breaker significantly reduced collagen gel contraction in immortalized human lung adenocarcinoma–derived CAFs (Fig. 4C-D).

We examined the expression of key collagen-modifying enzymes to determine whether defects in collagen compaction were attributable not only to impaired actomyosin-driven force generation but also to altered enzymatic maturation and crosslinking. Quantitative PCR analysis revealed downregulation of *lysyl oxidase* (*Lox*) and *procollagen-lysine,2-oxoglutarate 5-dioxygenase 1* (*Plod1*) in RAS–PI3K-deficient fibroblasts stimulated with KPB6-CM (Fig. 4E). Reintroduction of functional p110α restored the expression of both enzymes (Fig. 4E). Consistent with these findings, analysis of RNA from whole tumors in the *Pik3ca*^fRBD^ model revealed a significant reduction in *Lox* and *Plod1* expression compared with WT controls (Supplementary Fig. S4B). These data indicate that RAS–PI3K signaling regulates collagen maturation-associated machinery, extending its role beyond contractile force generation.

We next assessed *de novo* CAF-derived extracellular matrix (CDM) production using a glutaraldehyde crosslinked gelatin-based culture system^30^. Comparable cell viability and persistence were observed in WT and RBD fibroblast cultures throughout the matrix-generation period (Supplementary Fig. S4C). Sirius Red staining of CDMs produced by WT and RBD CAFs demonstrated that RAS–PI3K-deficient fibroblasts generated matrices containing significantly shorter and thinner collagen fibers compared with WT-derived CDMs (Fig. 4F–H). These structural defects mirrored the reduced and disorganized collagen deposition observed in tumors lacking stromal RAS–PI3K signaling (Fig. 3E), suggesting that impaired collagen assembly is an intrinsic property of RAS–PI3K-deficient fibroblasts. Reintroduction of functional p110α restored collagen fiber length but only partially rescued fiber thickness, whereas pharmacological PI3Kα inhibition in WT fibroblasts recapitulated the formation of shorter fibers (Fig. 4F–H). To further test whether the collagen-remodeling defect was fibroblast-intrinsic and independent of tumor-derived signals, we implanted collagen plugs containing TGF-β–activated WT or RBD fibroblasts into the flanks of syngeneic mice. Retrieved plugs containing WT fibroblasts showed long, well-organized collagen fibers, whereas RAS–PI3K-deficient fibroblasts produced shorter and structurally disorganized collagen fibers (Fig. 4I), confirming that RAS–PI3K-deficient fibroblasts have an intrinsic defect in collagen organization *in vivo*.

In addition to collagen, fibronectin deposition was also markedly reduced in tumors lacking stromal RAS–PI3K signaling (Fig. 3F), suggesting broader defects in ECM assembly. We therefore examined fibronectin organization using the CDM system to quantify overall matrix thickness and fibronectin fiber alignment. RAS–PI3K-deficient fibroblasts produced significantly thinner matrices with reduced fiber alignment compared to WT controls (Fig. 4J-L). Reintroduction of p110α restored fiber alignment but failed to fully rescue matrix thickness (Fig. 4J–L). Conversely, pharmacological inhibition of PI3Kα with alpelisib or disruption of RAS–PI3K coupling with the Breaker in WT murine CAFs phenocopied the disorganized and thinner matrix architecture observed in RAS–PI3K-deficient fibroblasts (Fig. 4J–L). Similar alterations were observed in human CAF-derived matrices following treatment with alpelisib or the RAS-PI3K Breaker (Supplementary Fig. S4D–E), supporting conservation of this RAS–PI3K-dependent matrix program across species. These matrix defects were also preserved under tumor-relevant conditions: when WT or RAS–PI3K-deficient CAFs were co-cultured with two distinct KRAS-mutant lung cancer cell lines, KPB6 and LKR13, RBD fibroblast-derived matrices displayed reduced fibronectin deposition and impaired fibrillar organization compared with WT-derived matrices (Supplementary Fig. S4F). Together, these findings are consistent with the stromal alterations observed in the *Pik3ca*^fRBD/-^ model (Fig. 3) and further demonstrate that RAS–PI3K signaling in CAFs governs not only ECM deposition but also higher-order matrix organization in tumor-associated contexts.

Given the pronounced alterations in collagen and fibronectin organization, we next examined whether RAS–PI3K disruption also affects the mechanical properties of CAF-derived matrices. CAF-derived matrices generated by RAS–PI3K-deficient fibroblasts exhibited a significantly lower Young’s modulus compared to WT matrices (Fig. 4M), indicating reduced matrix stiffness. Reintroduction of functional p110α partially restored matrix stiffness, whereas pharmacological inhibition of PI3Kα with alpelisib in WT fibroblasts phenocopied the mechanically softened RBD state (Fig. 4M).

To further examine matrix microstructural organization and molecular accessibility, we performed fluorescence lifetime imaging microscopy (FLIM) using the molecular rotor DCVJ, whose fluorescence lifetime increases when rotational motion becomes sterically restricted within dense or highly organized environments^57,58^. WT and RBD CAF-derived matrices were incubated with increasing concentrations of DCVJ and analyzed by phasor-based FLIM. While WT and RBD matrices displayed comparable fluorescence lifetimes at low dye concentrations, RBD-derived fibers exhibited significantly shorter lifetimes at higher concentrations, highlighted by blue-colored regions (Fig. 4N), indicating reduced steric restriction and altered microstructural organization. Phasor analysis revealed two major fluorescence lifetime populations centered around 0.3 ns and 1.9 ns, corresponding to freely diffusing and matrix-associated DCVJ molecules, respectively^59^. Based on this two-state model^60^, we estimated the fraction of dye associated with the matrix at increasing probe concentrations and observed a markedly reduced bound fraction in RBD-derived matrices compared to WT (Fig. 4O). Quantitative fitting of these data using a linear model based on phasor distributions indicated that WT matrices exhibited a higher apparent binding capacity than RBD matrices (Fig. 4O, inset). Consistent with the reduced bound fraction observed across DCVJ concentrations, these analyses indicate that RBD-derived matrices present fewer accessible probe-binding environments and altered microstructural organization compared with WT-derived matrices. Together, these findings indicate that WT-derived ECM fibers provide a more sterically restricted and molecularly accessible microenvironment for DCVJ binding, whereas RBD-derived matrices display reduced molecular restriction and altered microstructural organization.

### RAS–PI3K disruption remodels the CAF-derived matrisome and glycosylated ECM compartment

Having established that RAS–PI3K-deficient fibroblasts generate structurally disorganized, mechanically softer and microstructurally altered matrices, we next asked whether these defects were accompanied by broader changes in ECM composition. To address this, we performed quantitative proteomic analysis of decellularized CAF-derived matrices followed by matrisome-focused annotation.

Quantitative mass spectrometry identified widespread remodeling of the ECM proteome, with 515 differentially expressed proteins between WT and RAS–PI3K-deficient matrices. Further categorization of ECM components into core matrisome proteins (proteoglycans, ECM glycoproteins, and collagens) and matrisome-associated proteins (secreted factors, ECM-affiliated proteins, and ECM regulators) revealed 35 significantly deregulated matrisome-associated proteins, encompassing both core structural elements and matrisome-associated factors (Fig. 5A–B; Supplementary Fig. S5A and Supplementary Table S5). ECM glycoproteins represented the largest fraction of deregulated matrisome components, accounting for approximately 40% of all significant changes (Fig. 5C). Notably, whereas collagens and ECM-affiliated proteins were more frequently increased in RAS–PI3K-deficient matrices, glycoproteins were preferentially reduced (Supplementary Fig. S5B). This pattern suggests that RAS–PI3K disruption does not simply reduce global ECM abundance, but instead induces an imbalanced matrisome state in which structural collagen-associated components are uncoupled from glycoproteins involved in matrix organization, fibrillar connectivity and cell–matrix interactions. Consistently, functional enrichment analysis linked downregulated proteins to cell–matrix adhesion, TGF-β signaling and protein interaction networks, whereas upregulated components were associated with structural collagen pathways (Supplementary Fig. S5C). Importantly, this compositional imbalance provides a mechanistic framework for the architectural and mechanical defects observed in RAS–PI3K-deficient matrices, linking altered ECM composition to defective fibrillar organization and reduced matrix competence.

**Figure 5.**
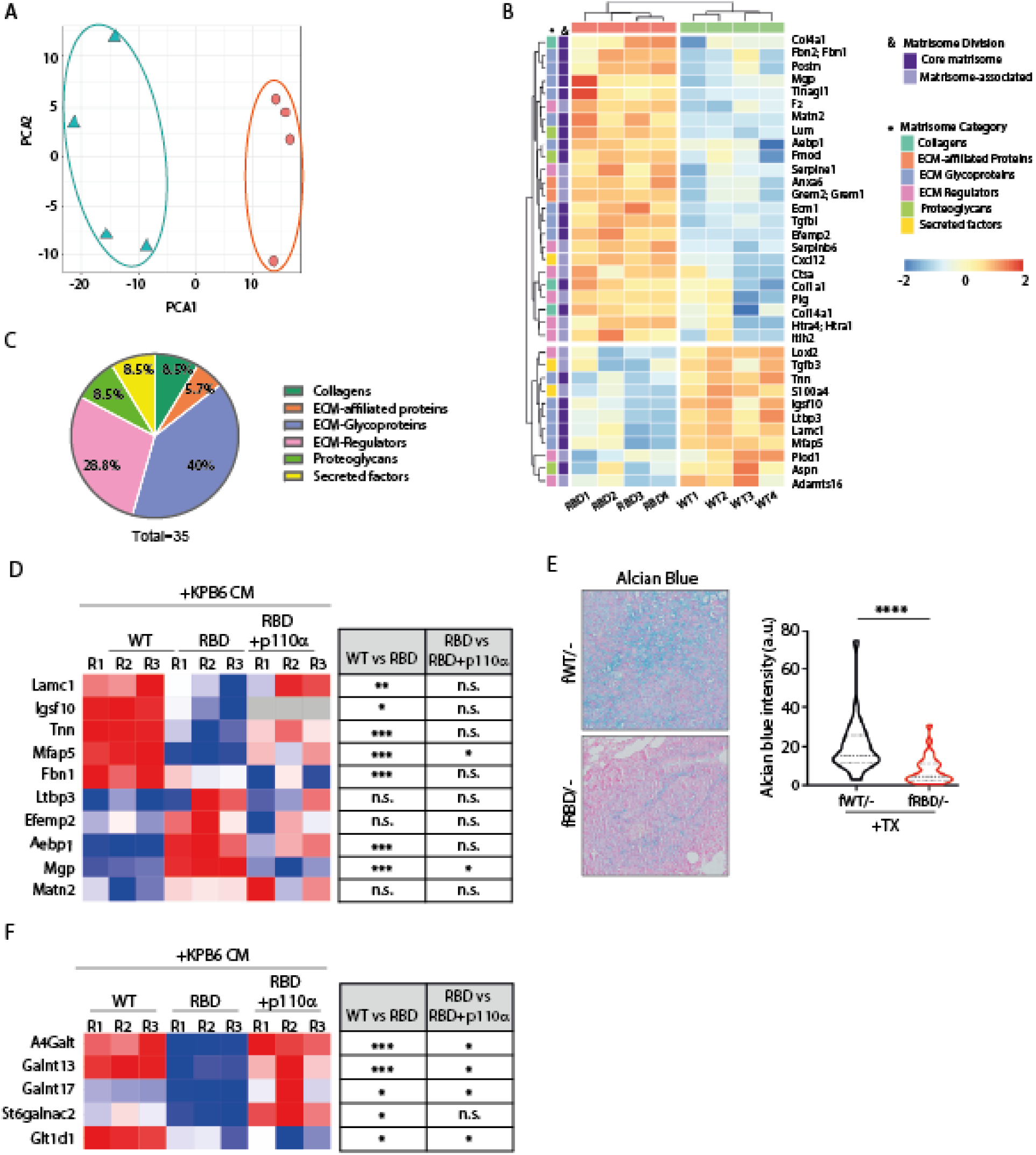
RAS–PI3K disruption remodels the CAF-derived matrisome and impairs the glycosylated ECM compartment. (A) Principal component analysis of quantitative proteomic profiles from decellularized CAF-derived matrices generated by WT and RAS–PI3K-deficient RBD fibroblasts, showing separation of WT- and RBD-derived matrisomes. (B) Heatmap of differentially regulated matrisome-associated proteins identified in WT and RBD CAF-derived matrices. Proteins are annotated according to matrisome division and category, including core matrisome and matrisome-associated components. Each column represents an independent biological replicate. (C) Pie chart showing the distribution of significantly deregulated matrisome components by category in RBD versus WT CAF-derived matrices. ECM glycoproteins represent the largest deregulated class. (D) Heatmap showing quantitative PCR validation of selected deregulated ECM glycoprotein-associated genes in WT, RBD and RBD+p110α fibroblasts following stimulation with KPB6-CM. Statistical comparisons for WT versus RBD and RBD versus RBD+p110α are shown on the right. Each column represents an independent biological replicate. (E) Representative Alcian Blue staining and quantification in tumors from tamoxifen-treated *Pik3ca*^fWT/−^ and *Pik3ca*^fRBD/−^ mice, showing reduced accumulation of acidic carbohydrate-rich extracellular components upon CAF-specific RAS–PI3K disruption. Data are shown as violin plots with median and quartiles. (F) Heatmap showing quantitative PCR validation of glycosyltransferases identified as deregulated in the RNA-seq analysis of WT, RBD and RBD+p110α fibroblasts stimulated with KPB6-CM. Statistical comparisons for WT versus RBD and RBD versus RBD+p110α are shown on the right. Each column represents an independent biological replicate. Statistical significance was determined using Student’s t-test unless otherwise indicated. *p < 0.05; **p < 0.01; ***p < 0.001; ****p < 0.0001; n.s., not significant.

To determine whether these matrisome alterations reflected CAF-intrinsic changes in ECM gene expression, we analyzed glycoproteins identified in the proteomic dataset by quantitative PCR. Several structural and matrix-associated glycoprotein genes, including *Lamc1*, *Igsf10*, *Tnn*, *Mfap5* and *Fbn1*, were reduced in RAS–PI3K-deficient fibroblasts, whereas *Aebp1* and *Mgp* were increased (Fig. 5D). Thus, RAS–PI3K disruption does not globally suppress glycoprotein expression, but reshapes the glycoprotein program produced by CAFs. Re-expression of functional p110α partially restored selected deregulated glycoproteins, including *Mfap5* and *Mgp*, supporting a direct contribution of RAS–PI3K signaling to this transcriptional program.

To determine whether alterations in the glycosylated ECM compartment are preserved *in vivo*, we analyzed lung tumors from fibroblast-specific RAS–PI3K-deficient mice using Alcian Blue staining. Alcian Blue detects acidic carbohydrate-rich extracellular components, including glycosaminoglycans and acidic glycoconjugates, thereby providing a histochemical readout of the glycosylated matrix compartment. Tumors from *Pik3ca*^fRBD/-^ mice showed a marked reduction in Alcian Blue staining compared with *Pik3ca*^fWT/-^ tumors (Fig. 5E), indicating that disruption of stromal RAS–PI3K signaling is associated with reduced accumulation of acidic carbohydrate-rich ECM components in vivo.

Because incorporation and maturation of glycoproteins within the ECM require post-translational processing, we next interrogated our RNA-seq dataset to identify glycosyltransferases deregulated upon RAS–PI3K disruption. *A4GALT*, *GALNT13*, *GALNT17*, *GLT1D1* and *ST6GALNAC2* were significantly downregulated in RBD CAFs stimulated with KPB6-CM. Quantitative PCR analysis confirmed reduced expression of these enzymes in RAS–PI3K-deficient fibroblasts compared with controls, whereas re-expression of functional p110α restored their levels in all cases except *GLT1D1* (Fig. 5F), indicating that their regulation is largely dependent on RAS–PI3K signaling. Consistent with the direction of expression changes observed in RBD CAFs, several RAS–PI3K-regulated glycoproteins and glycosyltransferases were associated with patient outcome in lung adenocarcinoma datasets (Supplementary Fig. S5D–E), supporting the clinical relevance of this ECM-associated program.

These findings indicate that RAS–PI3K signaling coordinates ECM assembly across structural, mechanical, and biochemical layers, thereby defining the matrix competence of CAFs and enabling the generation of organized and functionally mature extracellular matrices.

### RAS–PI3K-dependent ECM architecture controls tumor cell organization and growth

Having established that RAS–PI3K disruption alters CAF-derived ECM architecture, composition and mechanics, we next asked whether these changes influence how tumor cells interpret stromal matrix cues. To test this, we seeded tumor cells onto decellularized matrices generated by WT or RAS–PI3K-deficient CAFs. Tumor cells cultured on WT CAF-derived matrices aligned their actin cytoskeleton along the underlying fibronectin fibers, indicating that organized CAF-derived ECM provides directional cues for tumor cell organization (Fig. 6A). In contrast, RBD-derived matrices were less ordered and provided disorganized topographical cues, resulting in reduced and less coordinated tumor cell alignment (Fig. 6A). Quantitative analysis confirmed a strong correlation between fibronectin fiber orientation and tumor cell cytoskeletal alignment, consistent with ECM-mediated contact guidance (Fig. 6B). Similar matrix-guided organization was observed in an independent murine lung cancer cell line, LKR13 (Supplementary Fig. S6A–B), and in human lung cancer cells cultured on human CAF-derived matrices (Supplementary Fig. S6C–D). Pharmacological inhibition of PI3Kα in human CAFs similarly disrupted matrix organization and impaired directional tumor cell alignment (Supplementary Fig. S6C–D), indicating that RAS–PI3K-dependent control of matrix-guided tumor cell organization is conserved across species and pharmacologically targetable.

**Figure 6.**
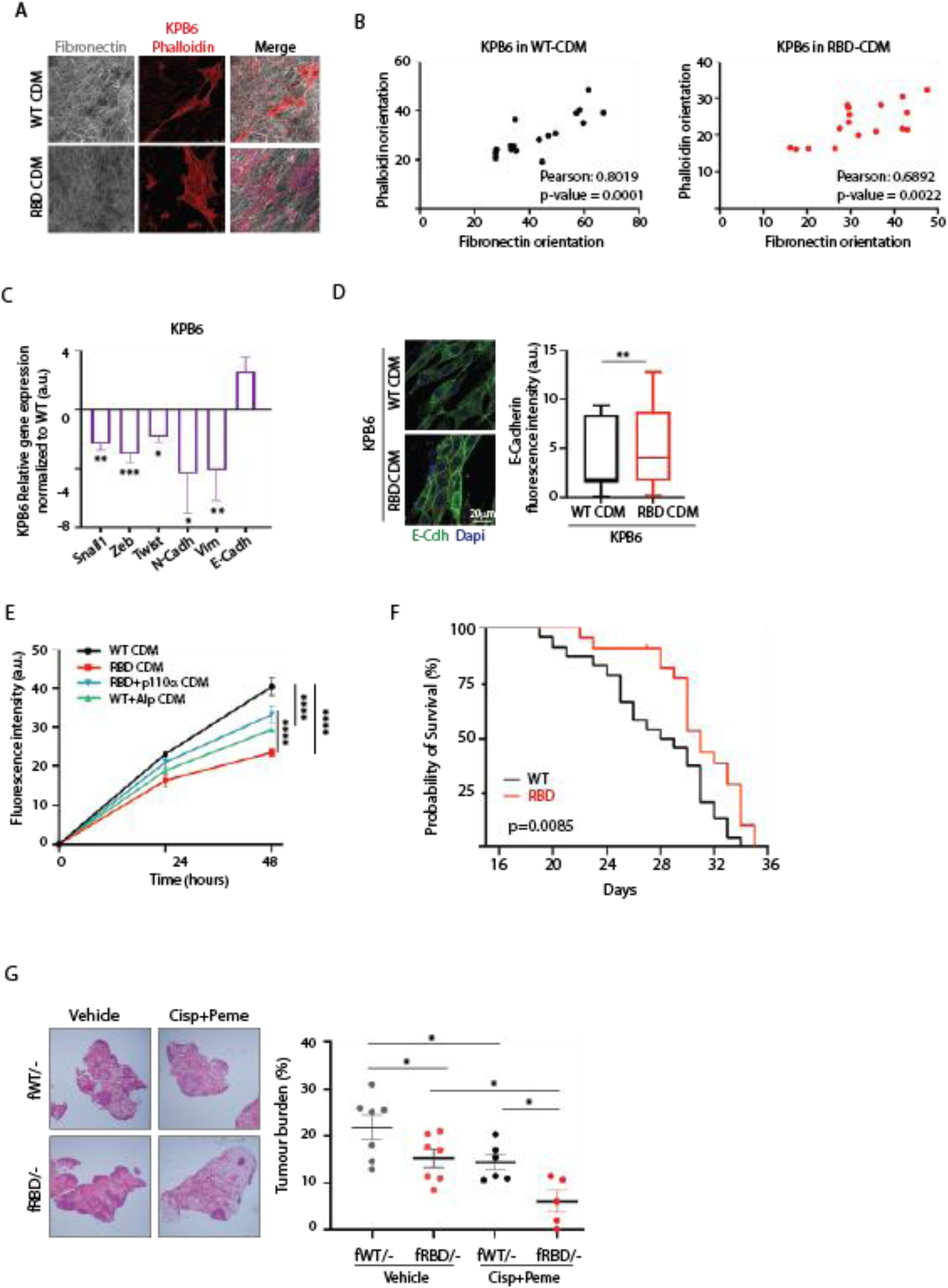
RAS–PI3K-dependent CAF-derived matrices provide directional and proliferative cues that support tumor cell progression and therapy resistance. (A) Representative immunofluorescence images of KPB6 tumor cells seeded onto decellularized CAF-derived matrices generated by WT or RBD fibroblasts. Fibronectin fibers are shown in grey and KPB6 actin cytoskeleton by phalloidin staining in red. Merged images show the relationship between tumor cell orientation and the underlying fibronectin network. (B) Correlation analysis between fibronectin fiber orientation and KPB6 phalloidin orientation in WT-and RBD-derived matrices. Pearson correlation coefficients and p values are indicated. (C) Quantitative PCR analysis of EMT-associated transcription factors and epithelial/mesenchymal markers in KPB6 cells cultured on RBD-derived matrices relative to WT-derived matrices. (D) Representative immunofluorescence images and quantification of E-cadherin expression in KPB6 cells cultured on WT- or RBD-derived matrices. E-cadherin is shown in green and nuclei in blue. Scale bar, 20 μm. (E) Proliferation analysis of KPB6 cells cultured on matrices generated by WT, RBD, RBD+p110α or alpelisib-treated WT fibroblasts, showing reduced tumor cell proliferation on RBD-derived matrices, partial rescue upon p110α re-expression, and phenocopying by pharmacological PI3Kα inhibition. (F) Kaplan–Meier survival analysis of *Pik3ca*^fWT/−^ and *Pik3ca*^fRBD/−^ mice following tamoxifen-induced recombination and intravenous injection of KPB6 lung cancer cells. (G) Representative H&E-stained lung sections and quantification of tumor burden in tamoxifen-treated *Pik3ca*^fWT/−^ and *Pik3ca*^fRBD/−^ mice treated with vehicle or cisplatin plus pemetrexed. Tumor burden is expressed as tumor area relative to total lung area. Each dot represents an individual mouse. Data are shown as mean ± SEM, box-and-whisker plots or individual values as indicated. Statistical significance was determined using Student’s t-test unless otherwise indicated, except for survival analysis, which was assessed using the log-rank test. *p < 0.05; **p < 0.01; ***p < 0.001; ****p < 0.0001.

We next examined whether the altered architecture of RBD-derived matrices affects tumor cell–matrix coupling. Tumor cells cultured on WT matrices formed focal adhesions that were strongly aligned with and positioned along fibronectin fibers. In contrast, cells seeded on RBD-derived matrices displayed reduced focal adhesion alignment and diminished colocalization with ECM fibers (Supplementary Fig. S6E–G). Re-expression of p110α in RBD CAFs restored focal adhesion–fiber coupling, whereas pharmacological inhibition of PI3Kα in WT CAFs impaired this interaction (Supplementary Fig. S6E–G). These findings indicate that RAS–PI3K-dependent ECM organization provides adhesive and topographical cues required for efficient tumor cell–matrix coupling.

We next asked whether altered ECM organization and tumor cell–matrix coupling influenced tumor cell phenotypic states. KPB6 cells cultured on RBD-derived matrices displayed reduced expression of EMT-associated transcription factors, including *Snail1*, *Zeb* and *Twist*, together with lower expression of mesenchymal markers such as N-cadherin (*Cdh2*) and vimentin (*Vim*), compared with cells grown on WT matrices (Fig. 6C). Conversely, expression of the epithelial marker *E-cadherin* was increased in tumor cells cultured on RBD-derived matrices, a finding further confirmed by immunofluorescence (Fig. 6D). Similar changes in EMT-associated gene expression were observed in LKR13 cells cultured on RBD-derived matrices (Supplementary Fig. S6H).

Consistent with the changes observed in tumor cell organization and EMT-associated programs, KPB6 cells cultured on matrices produced by RBD CAFs exhibited significantly reduced proliferation compared with cells grown on WT matrices (Fig. 6E). This defect was partially rescued by matrices generated by RBD CAFs re-expressing p110α, whereas pharmacological inhibition of PI3Kα in WT CAFs reduced the proliferative capacity of their matrices, phenocopying the RBD condition (Fig. 6E). Similar results were obtained with LKR13 cells (Supplementary Fig. S6I) and with human tumor cells cultured on human CAF-derived matrices (Supplementary Fig. S6J), indicating that the pro-proliferative effects of RAS–PI3K-dependent CAF-derived ECM are conserved across tumor models.

These findings provide a mechanistic framework for the reduced tumor burden and decreased tumor cell proliferation observed in the *Pik3ca*^fRBD/-^ model (Fig. 3), linking stromal RAS–PI3K signaling and ECM organization to tumor growth control. In line with this, *Pik3ca*^fRBD/-^ mice displayed significantly prolonged survival compared with WT animals (Fig. 6F). Notably, mortality onset was markedly delayed in *Pik3ca*^fRBD/-^ mice, suggesting that CAF-specific disruption of RAS–PI3K signaling generates a less permissive tumor microenvironment that slows tumor progression *in vivo*.

Because disruption of stromal RAS–PI3K signaling reshaped the tumor microenvironment and limited tumor progression, we next asked whether this altered stromal state affects therapeutic response. *Pik3ca*^fRBD/-^ mice displayed significantly improved responses to cisplatin plus pemetrexed compared with WT controls (Fig. 6G), indicating that stromal RAS–PI3K signaling contributes to chemotherapy resistance.

Together, our data indicate that stromal RAS–PI3K signaling sustains a tumor-supportive CAF state that promotes ECM organization, tumor cell–matrix coupling, proliferative growth and chemotherapy resistance. These broad functional alterations, together with the inflammatory transcriptional rewiring identified in RAS–PI3K-deficient CAFs, prompted us to further examine the immunomodulatory programs engaged upon RAS–PI3K disruption.

### RAS–PI3K loss engages an IL6–STAT3-associated immunomodulatory program in CAFs

Transcriptomic analyses revealed that RAS–PI3K-deficient CAFs were enriched in inflammatory and immune-related programs, particularly in response to tumor-derived stimulation (Fig. 2). Further GSEA identified significant enrichment of IL6-associated signaling pathways in RBD CAFs, including the BIOCARTA_IL6_PATHWAY signature (Fig. 7A) and the REACTOME_INTERLEUKIN_6_FAMILY_SIGNALING pathway (Supplementary Fig. S7A), suggesting engagement of cytokine-associated inflammatory programs upon loss of RAS–PI3K signaling.

**Figure 7.**
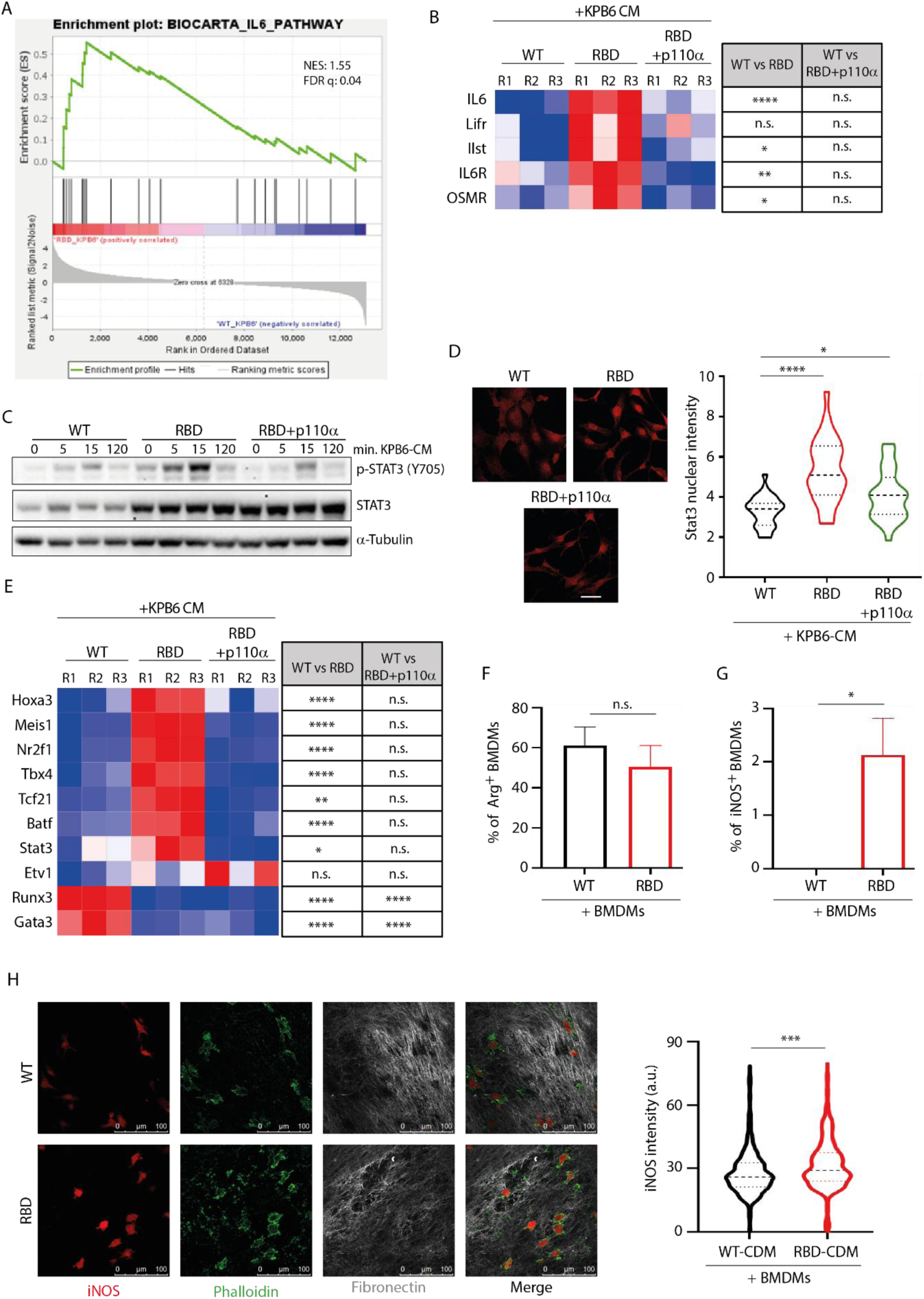
RAS–PI3K disruption engages an IL6–STAT3-associated immunomodulatory CAF program. (A) Gene Set Enrichment Analysis showing enrichment of the BIOCARTA_IL6_PATHWAY signature in RBD CAFs compared with WT CAFs following stimulation with KPB6-CM. Normalized enrichment score (NES) and false discovery rate (FDR q value) are indicated. (B) Heatmap showing quantitative PCR analysis of IL6 signaling-associated genes, including Il6 and components of the IL6 receptor complex, in WT, RBD and RBD+p110α fibroblasts stimulated with KPB6-CM. Statistical comparisons for WT versus RBD and WT versus RBD+p110α are shown on the right. Each column represents an independent biological replicate. (C) Immunoblot analysis of STAT3 phosphorylation at Y705 and total STAT3 levels in WT, RBD and RBD+p110α fibroblasts stimulated with KPB6-CM for the indicated times. α-Tubulin was used as loading control. (D) Representative immunofluorescence images and quantification of nuclear STAT3 accumulation in WT, RBD and RBD+p110α fibroblasts following stimulation with KPB6-CM, showing increased STAT3 nuclear localization in RBD fibroblasts and reduction upon p110α re-expression. (E) Heatmap showing quantitative PCR analysis of selected transcriptional regulators associated with fibroblast plasticity and immunomodulatory stromal programs in WT, RBD and RBD+p110α fibroblasts stimulated with KPB6-CM. Statistical comparisons for WT versus RBD and WT versus RBD+p110α are shown on the right. Each column represents an independent biological replicate. (F) Flow cytometry analysis of arginase expression in bone marrow-derived macrophages (BMDMs) co-cultured with WT or RBD CAFs for 24 h. (G) Flow cytometry analysis of iNOS expression in BMDMs co-cultured with WT or RBD CAFs for 24 h, showing emergence of an iNOS-positive macrophage population upon co-culture with RBD CAFs. (H) Representative immunofluorescence images and quantification of BMDMs seeded onto decellularized CAF-derived matrices generated by WT or RBD fibroblasts. BMDMs were stained for iNOS and phalloidin, and fibronectin staining was used to visualize the underlying CAF-derived matrix. Quantification shows increased iNOS fluorescence intensity in macrophages seeded onto RBD-derived matrices compared with WT-derived matrices. Data are shown as mean ± SEM or violin plots with median and quartiles, as indicated. Statistical significance was determined using Student’s t-test unless otherwise indicated. *p < 0.05; **p < 0.01; ***p < 0.001; ****p < 0.0001; n.s., not significant.

Consistent with this, *Il6* expression was markedly increased in RBD CAFs compared with WT controls following stimulation with tumor-conditioned medium, and this increase was reversed by re-expression of functional p110α (Fig. 7B), indicating that *IL6* induction is linked to disruption of the RAS–PI3K axis. RBD CAFs also showed increased expression of components of the *IL6* receptor complex, including *Il6r*, *Osmr*, *Il6st* and *Lifr*, by quantitative PCR (Fig. 7B). These data indicate that RAS–PI3K-deficient CAFs engage both ligand and receptor components of an IL6-associated signaling module.

We next examined whether this IL6-associated program was accompanied by activation of STAT3 signaling. Following stimulation with KPB6-CM, RBD CAFs displayed increased total STAT3 and enhanced phosphorylation of STAT3 at Y705 compared with WT controls (Fig. 7C and Supplementary Fig. S7B). Re-expression of functional p110α markedly reduced STAT3 phosphorylation, although total STAT3 protein levels remained elevated in RBD+p110α cells. Consistent with pathway activation, immunofluorescence analysis revealed increased nuclear accumulation of STAT3 in RBD CAFs compared with WT controls, both at early (15 min) (Supplementary Fig. S7C) and sustained (24 h) time points (Fig. 7D) following stimulation with KPB6-CM. Re-expression of functional p110α reduced nuclear STAT3 accumulation under both conditions (Fig. 7D and Supplementary Fig. S7C), further supporting activation of a persistent STAT3-associated signaling state upon loss of RAS–PI3K signaling.

To determine whether STAT3 pathway activation was engaged within a broader regulatory program, we examined transcriptional regulators differentially expressed in our RNA-seq dataset and associated with fibroblast plasticity or immunomodulatory stromal programs^61,62^. RAS–PI3K-deficient CAFs showed increased expression of several regulators, including *Hoxa3*, *Meis1*, *Nr2f1*, *Tbx4*, *Tcf21* and *Batf*, most of which were restored toward WT levels upon re-expression of functional p110α (Fig. 7E). In contrast, *Runx3* and *Gata3* were reduced in RBD CAFs and were not rescued by p110α re-expression.

Additional secretory and fibroblast-state markers, including *Cxcl12* and *Pdgfrα*, were also altered in RBD CAFs and partially normalized upon p110α re-expression (Supplementary Fig. S7D–E), consistent with broader remodeling of the CAF secretory and regulatory landscape beyond the IL6–STAT3 axis.

To determine whether the immunomodulatory program induced by RAS–PI3K loss in CAFs could influence macrophage state, we co-cultured bone marrow-derived macrophages (BMDMs) with WT or RBD CAFs and analyzed macrophage polarization-associated markers by flow cytometry. Arginase expression tended to be lower in BMDMs co-cultured with RBD CAFs compared with WT CAFs, although this difference did not reach statistical significance (Fig. 7F). In contrast, iNOS expression was negligible in BMDMs co-cultured with WT CAFs but became detectable in a distinct BMDM population after 24 h of co-culture with RBD CAFs (Fig. 7G), indicating that RAS–PI3K-deficient CAFs can promote engagement of an iNOS-associated macrophage state.

Because RAS–PI3K disruption profoundly remodels CAF-derived ECM architecture, we next asked whether the altered matrix itself could influence macrophage phenotype. BMDMs seeded onto decellularized RBD-derived matrices showed increased iNOS staining compared with BMDMs seeded onto WT-derived matrices (Fig. 7H).

Importantly, this immunomodulatory program emerged in a context in which fibroblast-specific RAS–PI3K disruption also altered tumor immune composition in vivo, reducing CD68⁺ macrophage accumulation and increasing CD8⁺ T cell infiltration (Fig. 3). Although the causal contribution of the STAT3-associated CAF program and the iNOS-associated macrophage response to immune remodeling in vivo remains to be determined, these data indicate that RAS–PI3K disruption does not simply impair myofibroblastic CAF activation, but promotes engagement of an alternative immunomodulatory stromal program with functional consequences for macrophage state.

## DISCUSSION

CAF plasticity is increasingly recognized as a major determinant of tumor progression^10,13^, yet the signaling mechanisms that stabilize specific CAF states remain incompletely understood. Our findings identify RAS–PI3K signaling as a central regulator of CAF functional identity in lung cancer. Rather than simply attenuating fibroblast activation, disruption of RAS–PI3K signaling redirected CAFs away from a myofibroblastic, matrix remodeling state and toward an immunomodulatory transcriptional program. This reprogramming was accompanied by profound defects in ECM assembly, altered tumor cell behavior, reduced tumor progression, and increased therapeutic vulnerability. Together, these data support a model in which stromal RAS–PI3K signaling functions as a regulatory node that coordinates CAF state, ECM competence and the tumor-supportive properties of the microenvironment.

The coexistence of myofibroblastic, inflammatory, antigen-presenting and other CAF states has been described across tumor types^3,4,7,63-66^, but how stromal cells interpret tumor-derived cues to adopt one functional state over another remains a key unresolved question. Our data suggest that RAS–PI3K signaling contributes to this decision by maintaining a matrix-producing myofibroblastic program while restraining inflammatory and immune-related transcriptional modules. Importantly, loss of RAS–PI3K signaling did not simply silence CAF activation. Instead, RAS–PI3K-deficient CAFs retained responsiveness to tumor-derived cues but interpreted these signals through a distinct transcriptional framework, marked by reduced matrix-associated gene expression and enrichment of immunomodulatory programs. This places RAS–PI3K signaling as a regulator of CAF state balance rather than a generic activator of fibroblast function. Our study therefore adds to current CAF plasticity frameworks by identifying a genetically defined stromal signaling axis that couples tumor-derived cue interpretation to the execution of a matrix-remodeling CAF state^6,11-13,67,68^.

Mechanistically, the earliest detectable consequence of RAS–PI3K disruption was a failure to engage the cytoskeletal remodeling program required for myofibroblastic CAF execution^28,69-71^. RAS–PI3K-deficient fibroblasts preserved canonical TGF-β signal perception, but failed to polymerize F-actin efficiently, organize stress fibers, and induce α-SMA and vimentin in response to profibrotic or tumor-derived cues.

This distinction suggests that RAS–PI3K signaling does not simply regulate upstream responsiveness to activating stimuli, but rather controls the ability of CAFs to translate these cues into a contractile and matrix-remodeling phenotype. This is particularly relevant because actin cytoskeletal remodeling is tightly linked to fibroblast mechanotransduction: stress fibers and focal adhesions enable cells to generate traction forces, sense matrix stiffness and mechanically remodel collagen- and fibronectin-rich matrices^5,28,72,73^. Through these processes, cytoskeletal tension can reinforce ECM organization and sustain matrix-producing myofibroblastic programs^5,28,29,70,74-79^. Thus, RAS–PI3K signaling may stabilize myCAF identity by coupling biochemical activation signals to the cytoskeletal machinery required for contractile force generation, matrix remodeling, and functional ECM assembly. When this node is disrupted, CAFs fail to consolidate a myofibroblastic state and become permissive to alternative immunomodulatory transcriptional trajectories.

Downstream of this impaired cytoskeletal and contractile execution, a major consequence of RAS–PI3K loss was a failure of matrix assembly competence. RAS–PI3K-deficient CAFs did not merely produce less organized collagen and fibronectin networks; they generated matrices with altered mechanical, microstructural and compositional properties. Matrisome analysis further revealed broader ECM remodeling, particularly affecting ECM glycoproteins and glycosylation-associated enzymes, suggesting that RAS–PI3K signaling coordinates matrix competence at multiple levels, from cytoskeletal force generation to ECM maturation and higher-order organization. These findings are consistent with the current view of ECM assembly as a multistep process in which composition, post-translational modification, protein–protein interactions and cell-mediated remodeling converge to generate functionally specialized matrix architectures. Thus, the matrix phenotype induced by RAS–PI3K loss cannot be explained by changes in individual ECM components alone. Instead, RAS–PI3K signaling appears to integrate cytoskeletal, biochemical and post-translational layers of ECM assembly to generate a functionally competent stromal matrix^32^. Given the central role of ECM architecture and mechanics in regulating tumor cell behavior^16,80-83^, therapeutic response^25,84,85^ and immune cell localization^24-26,86,87^, these results identify RAS–PI3K signaling as a stromal mechanism controlling the instructive properties of the tumor ECM.

The functional relevance of this loss of matrix competence was evident in tumor cells cultured on CAF-derived matrices. A competent stromal matrix not only provides structural support, but also delivers directional, adhesive and mechanical cues that instruct tumor cell behavior^26,80,88,89^. WT CAF-derived matrices promoted tumor cell alignment, focal adhesion coupling, EMT-associated gene expression and proliferative growth, whereas matrices generated by RAS–PI3K-deficient CAFs failed to efficiently support these responses. These observations suggest that RAS–PI3K-dependent ECM assembly generates a tumor-supportive matrix niche capable of reinforcing malignant behavior. In line with this, CAF-specific disruption of RAS–PI3K signaling reduced tumor burden, decreased p-Histone H3 staining and prolonged survival *in vivo*. The increased sensitivity of tumor cells grown on RBD-derived matrices to cisplatin plus pemetrexed, together with the improved therapeutic response observed in *Pik3ca*^fRBD/-^tumors, further suggests that stromal matrix competence contributes to microenvironmental protection from chemotherapy^84,90^. Thus, RAS–PI3K signaling enables CAFs to build an instructive matrix niche that supports tumor growth and limits therapeutic efficacy, at least in part by preserving the adhesive, directional and mechanical cues required for tumor cell–ECM coupling.

In parallel with ECM remodeling, RAS–PI3K disruption promoted a STAT3-linked immunomodulatory CAF program. RAS–PI3K-deficient CAFs showed increased expression of *IL6*, *CXCL12*, *PDGFRα* and transcriptional regulators associated with fibroblast plasticity and immune programs, together with enhanced STAT3 activation and nuclear accumulation. Importantly, this program was functionally linked to macrophage state. In co-culture experiments, BMDMs exposed to RAS–PI3K-deficient CAFs showed negligible changes in arginase expression, but acquired a detectable iNOS-positive population that was largely absent in co-cultures with WT CAFs. Moreover, BMDMs seeded onto decellularized RBD-derived matrices displayed increased iNOS expression compared with those cultured on WT-derived matrices. These findings indicate that RAS–PI3K loss in CAFs can influence macrophage phenotype through both cellular crosstalk and matrix-dependent mechanisms. *In vivo*, CAF-specific RAS–PI3K disruption was associated with reduced CD68⁺ macrophage accumulation and increased CD8⁺ T cell infiltration, suggesting that the stromal state induced by RAS–PI3K loss is associated with a remodeled immune contexture. Importantly, this immunomodulatory program should not be interpreted as intrinsically tumor-suppressive. Rather, in the lung cancer models studied here, it emerged together with loss of matrix competence, induction of iNOS-associated macrophage features, altered immune composition and impaired tumor progression. This pattern is compatible with a less tumor-permissive immune contexture^91-93^, although the functional state and causal contribution of these immune populations remain to be established.

A key remaining question is how the matrix-remodeling and immunomodulatory arms of the RAS–PI3K-deficient CAF state are mechanistically connected. Recent ecosystem-level models of the TME emphasize that ECM architecture is not merely structural, but acts as an organizing scaffold that determines where immune cells localize and how they behave ^7,26^. In this framework, fibroblast-derived ECM programs establish permissive or restrictive immune niches, while ECM composition and mechanics can directly influence macrophage and T cell behaviour ^4,7,94-96^. Our data support this concept by showing that CAF-specific RAS–PI3K disruption simultaneously impairs matrix assembly, alters immune composition *in vivo* and generates matrices that directly promote increased iNOS expression and morphological changes in macrophages. These findings suggest that defective matrix competence may contribute to immune remodeling, although soluble CAF-derived mediators are also likely to participate. In addition, multiomic analyses have identified conserved mechanoresponsive and immunomodulatory CAF states^6^, supporting the idea that mechanical and immune-related fibroblast programs are functionally connected rather than fully independent. One potential link between these programs may involve mechanotransduction-dependent transcriptional regulators. YAP/TAZ activity is known to respond to cytoskeletal tension and matrix mechanics, and recent work in prostate cancer showed that YAP1 sustains ECM-producing CAF states while restraining NF-κB-associated lymphocyte-activating CAF programs^97^. Although YAP/TAZ activity was not directly tested as a causal mediator in this study, these observations raise the possibility that loss of RAS–PI3K-dependent cytoskeletal and matrix competence weakens mechanotransductive programs that normally stabilize myCAF identity, thereby facilitating engagement of STAT3/NF-κB-linked immunomodulatory programs. Whether the immune remodeling observed in *Pik3ca*^fRBD/-^ tumors is driven by altered matrix architecture, changes in CAF-derived soluble factors, reduced tumor burden, or a combination of these mechanisms remains to be determined.

Importantly, our genetic rescue and pharmacological inhibition experiments indicate that the stromal program controlled by RAS–PI3K signaling is pharmacologically modifiable. Re-expression of functional p110α restored multiple defects observed in RAS–PI3K-deficient CAFs, supporting a direct role for this signaling axis in regulating matrix assembly. Conversely, PI3Kα inhibition with alpelisib recapitulated key features of the RAS–PI3K-deficient ECM phenotype, including impaired collagen contraction, altered fibronectin organization and reduced matrix stiffness. Pharmacological disruption of RAS–PI3K coupling with the Breaker also impaired collagen contraction and fibronectin organization in both murine and human CAFs, supporting the tractability of this stromal signaling axis. Together, these findings suggest that CAF matrix competence is not a fixed stromal property, but a pharmacologically modifiable state. This interpretation is consistent with recent work in colorectal cancer identifying PI3K/AKT/mTOR signalling as divergent regulators of CAF plasticity^98^. In that study, PI3K/mTOR inhibition reduced myCAF-associated programs and promoted inflammatory CAF features through an FGF2–FGFR1–JAK2–STAT3 axis, although this transition generated a protumorigenic inflammatory state. Together with our findings, these data support the idea that PI3K pathway activity contributes to myofibroblastic CAF programs, while highlighting that the functional outcome of PI3K pathway disruption is highly context-dependent. In our lung cancer models, selective disruption of RAS–PI3K coupling impaired ECM competence, reduced tumor growth, enhanced therapeutic vulnerability and was associated with reduced macrophage accumulation and increased CD8⁺ T cell infiltration. Thus, targeting stromal RAS–PI3K signaling should not be viewed as a generic CAF-inhibitory strategy, but as a context-dependent approach to reprogram specific tumor-supportive stromal functions.

Several aspects of this work warrant further investigation. First, although our genetic model enables fibroblast-specific disruption of RAS–PI3K signaling, CAFs are heterogeneous and it remains unclear whether all CAF subsets depend equally on this pathway. Single-cell and spatial approaches will be needed to determine whether RAS–PI3K disruption selectively prevents the emergence of particular myofibroblastic states, promotes defined immunomodulatory subsets, or induces a hybrid state not fully captured by canonical CAF classifications^7,11,13,67^. Second, the survival associations observed for selected glycoproteins and glycosylation-related enzymes in lung adenocarcinoma cohorts are correlative and based on bulk tumor datasets. These analyses support clinical relevance but do not establish CAF-specific prognostic function. Third, while the improved response to cisplatin plus pemetrexed suggests that RAS–PI3K-dependent matrices contribute to therapeutic resistance, additional studies will be required to determine whether this reflects altered drug penetration, reduced survival signaling, changes in tumor cell state, or broader remodeling of the microenvironment^26,83-85^. Finally, although CAF-specific RAS–PI3K disruption was associated with reduced macrophage accumulation and increased CD8⁺ T cell infiltration, the causal contribution of these immune populations to reduced tumor progression remains to be defined. Our macrophage co-culture and matrix-seeding experiments indicate that RAS–PI3K-deficient CAFs and their matrices can promote iNOS-associated macrophage features, but additional studies will be required to determine the full functional identity of these macrophages *in vivo*, their impact on T cell recruitment or activity, and whether they contribute directly to tumor control. In addition, our current data do not distinguish the relative contribution of CAF-derived soluble mediators, altered ECM architecture, reduced tumor burden, or combinations of these mechanisms to immune remodeling *in vivo*.

Overall, our study expands the role of RAS signaling in cancer beyond tumor cell–intrinsic oncogenic pathways by identifying stromal RAS–PI3K signaling as a regulator of CAF functional plasticity. By enabling CAFs to translate tumor-derived cues into cytoskeletal remodeling, matrix maturation and higher-order ECM organization, this pathway supports the formation of a tumor-permissive stromal niche. Its disruption uncouples CAFs from myofibroblastic matrix competence, permits engagement of alternative immunomodulatory programs, reshapes macrophage–matrix interactions and weakens the physical and cellular support required for tumor progression and chemotherapy resistance. These findings support a framework in which stromal therapies are designed not to eliminate CAFs indiscriminately, but to reprogram defined tumor-supportive CAF functions.

## ACKNOWLEDGEMENTS

This work was supported by grants from the Spanish Ministry of Science and Innovation (RTI2018-099161-A-I00, PID2024-162333OB-I00), Rosetrees Trust, AECC Excellence program Stop Ras Cancers (EPAEC222641CICS), Programa JAE-Intro ICU from CSIC (JAEICU-21-IBMCC-6) and JCyL (CSI185-20). This research was co-financed by FEDER funds. The CIC is supported by «Escalera de Excelencia» of the Education Ministry of the Castilla y León autonomous government plus the European Regional Development Fund (CLU-2023-2-01). The authors wish to thank the Pathology Unit, the Mouse Model Experimentation Unit, and the Advanced Cellular Analysis Unit at CIC for their assistance in carrying out this work. The authors thank members of the lung TRACERx consortium whose study derived the human cancer-associated fibroblasts (CAFs) that were used in this study. The TRACERx study is funded by Cancer Research UK (CRUK; C11496/A17786) and the creation of TRACERx patient-derived models was supported by the Wellcome Trust (209199/Z/17/Z), the CRUK Lung Cancer Centre of Excellence and the Francis Crick Institute, which receives its core funding from CRUK (FC001169), the MRC (FC001169), and the Wellcome Trust (FC001169). This work was supported in part by research funding from BridgeBio Oncology Therapeutics.

## MATERIALS AND METHODS

### Mouse model generation and experimental procedures

The mouse model with fibroblast-specific RAS–PI3K disruption was generated by crossing *Pik3ca^RBD/flox^* mice ^50^ with Tg(Col1a2-cre/ERT,-ALPP)7Cpd/J (The Jackson Laboratory; referred to here as *Col1a2-CreER*). For autochthonous tumor studies, this model was further bred with KRAS^LA2^ mice^54^.

For the experimental lung colonization model, 1x10^5^ KPB6 tumor cells were injected through the tail vein of *Pik3ca^fRBD/-^*mice in a total volume of 100 µL PBS. For tumor growth experiments, mice were sacrificed 21 days post-injection, and lungs were collected for histological analysis. For survival studies, mice were monitored daily and euthanized upon reaching humane endpoints, including a 20% reduction in body weight relative to baseline or signs of respiratory distress, according to approved ethical guidelines. For therapeutic studies, drug treatments began 1 week after tail vein injection.

All animals were housed and maintained at the NUCLEUS animal facility of the University of Salamanca under standard pathogen-free conditions. Animal care and experimental procedures were conducted in strict compliance with European (2007/526/CE) and Spanish (RD 1201/2005; RD 53/2013) legislation, and the study was approved by the Bioethics Committee of the Cancer Research Center.

Efforts were made to minimize animal suffering, and humane endpoints were established in accordance with the approved protocols. Both male and female mice were used in this study, and experimental groups were balanced for sex and birth date whenever possible. No overt sex-dependent differences were observed in the experimental outcomes analyzed. Animals were randomly assigned to experimental groups to ensure unbiased allocation. Researchers were blinded to group assignments during data collection and analysis to reduce bias. No formal sample size calculation was performed; however, sample sizes were determined based on prior experience and feasibility. Pre-established inclusion and exclusion criteria were applied to maintain experimental integrity. Animals presenting pre-existing health conditions were excluded.

### *In Vivo* Tamoxifen and Drug Treatments

Mice were treated with tamoxifen (3.2 mg, dissolved in 80 µL of corn oil; MedChemExpress) by oral gavage for 3 consecutive days. For long-term experiments, tamoxifen administration was repeated once every 2 weeks throughout the duration of the study to maintain Cre recombination efficiency.

For chemotherapy studies, cisplatin plus pemetrexed treatment was administered intraperitoneally twice a week for 2 consecutive weeks. Mice received either vehicle or combination treatment with cisplatin (3 mg/kg/day; MedChemExpress) and pemetrexed (100 mg/kg/day; MedChemExpress). Cisplatin and pemetrexed were dissolved in NaCl 0.9%.

### Tissue culture

Cells were cultured in Dulbecco’s Modified Eagle Medium (DMEM; Gibco) supplemented with 10% fetal bovine serum (FBS; Gibco), 100 U/mL penicillin, 100 µg/mL streptomycin (Pen/Strep; Gibco) and 1% glutamine. Cells were maintained in a humidified incubator at 37°C with 5% CO₂.

Murine embryonic fibroblasts (MEFs) were activated with TGF-β1 (10 ng/mL; PeproTech) for 24 h or treated with conditioned medium (CM) from the KRAS-mutant lung cancer cell lines KPB6 and LKR13. CM was prepared by culturing 1.5 × 10⁶ cancer cells in DMEM supplemented with 5% FBS for 48 h, followed by filtration through a 0.45 µm filter. Fibroblasts were incubated with CM for 24 h.

Human CAF (hCAF) cultures were explanted from tumor tissue from patients with lung adenocarcinoma within the Tracking Cancer Evolution through Therapy (TRACERx) clinical study (REC reference 13/LO/1546). hCAFs were activated similarly to murine CAFs using conditioned medium from A549 cells.

### *In vitro* pharmacological treatments

For pharmacological inhibition experiments, fibroblasts were treated with the PI3Kα inhibitor alpelisib (0.5 µM) or with the RAS–PI3K Breaker (1.0 µM), a small molecule that interferes with the RAS–PI3K interaction. Both compounds were dissolved in DMSO. Equivalent DMSO concentrations were used as vehicle controls. The same concentrations were used in murine and human fibroblast models unless otherwise indicated.

### CDM generation and decellularization

CDMs were generated and decellularized as previously described^99,100^. Briefly, fibroblasts were seeded onto gelatin-coated, glutaraldehyde-crosslinked culture dishes and maintained in matrix medium supplemented with ascorbic acid to promote ECM deposition.

For co-culture-derived CDMs, gelatin-coated plates were seeded with fibroblasts as described above, without prior activation with TGF-β1. The following day, 6 × 10⁵ KPB6 or LKR13 cancer cells were seeded onto the fibroblast monolayer. After 24 h, the culture medium was replaced with matrix medium consisting of DMEM supplemented with 50 µg/mL ascorbic acid. Medium was partially replaced every 48 h with fresh matrix medium containing 100 µg/mL ascorbic acid, thereby maintaining a final concentration of 50 µg/mL ascorbic acid throughout the culture period. After 8 days, non-decellularized CDMs were fixed for immunofluorescence analysis.

Where indicated, alpelisib (0.5 µM), the RAS-PI3K Breaker (1 µM) or an equivalent volume of DMSO vehicle was added to the matrix medium during CDM generation and refreshed with each medium change.

### Mass spectrometry

Proteomics experiments were performed using mass spectrometry as previously reported^101-103^ with some technical modifications. Peptides, digested from proteins using trypsin, were desalted with the AssayMAP Bravo (Agilent Technologies) platform using the peptide clean-up v3.0 protocol. Briefly, reverse phase S cartridges (Agilent, 5 μL bed volume) were primed with 250 μL 99.9% acetonitrile (ACN) with 0.1%TFA and equilibrated with 250 0.1% TFA at a flow rate of 10 μL/min. The samples were loaded at 20 μL/min, followed by an internal cartridge wash with 0.1% TFA at a flow rate of 10 μL/min. Peptides were then eluted with 105 μL of solution (70/30 ACN/ H2O + 0.1% TFA. Eluted peptide solutions were dried in a SpeedVac vacuum concentrator and peptide pellets were stored at −80 °C.Dried peptides were dissolved in 0.1% TFA and analysed by nanoflow ultimate 3000 RSL nano instrument was coupled on-line to a Q Exactive plus mass spectrometer (Thermo Fisher Scientific). Gradient elution was from 3% to 28% solvent B in 90 min at a flow rate 250 nL/min with solvent A being used to balance the mobile phase (buffer A was 0.1% formic acid in water and B was 0.1% formic acid in acetonitrile). The spray voltage was 1.95 kV and the capillary temperature was set to 255 °C. The Q-Exactive plus was operated in data dependent mode with one survey MS scan followed by 15 MS/MS scans. The full scans were acquired in the mass analyser at 375- 1500m/z with the resolution of 70 000, and the MS/MS scans were obtained with a resolution of 17 500. MS raw files were converted into Mascot Generic Format using Mascot Distiller (version 2.8.1) and searched against the SwissProt database (SwissProt_2021_02.fasta) restricted to human entries using the Mascot search daemon (version 2.8.0). Allowed mass windows were 10 ppm and 25 mmu for parent and fragment mass to charge values, respectively. Identified peptides were quantified using in-house software Pescal as described before^104,105^. The resulting quantitative data was parsed into R (Version 4.2.2) for further normalisation and statistical analysis. The code used in analysis and visualization of data is available at https://github.com/CutillasLab/protools2.

Phosphoproteomics experiments were performed using mass spectrometry as previously reported ^102,106^. In brief, frozen tissues were grounded and lysed in 8M urea buffer and supplemented with phosphatase inhibitors (10 mM Na_3_VO_4_, 100 mM β-glycerol phosphate and 25 mM Na_2_H_2_P_2_O_7_ (Sigma)). Proteins were digested into peptides as outlined above. Phosphopeptides were enriched from total peptides by TiO2 chromatography essentially as reported previously^107^. Dried phosphopeptides and peptides were dissolved in 0.1% TFA and analysed by LTQ-Orbitrap XL mass spectrometer (Thermo Fisher Scientific) connected to a nanoflow ultra-high-pressure liquid chromatography (UPLC, NanoAcquity, Waters). Peptides were separated using a 75 μm × 150 mm column (BEH130 C18, 1.7 μm Waters) using solvent A (0.1% FA in LC–MS grade water) and solvent B (0.1% FA in LC–MS grade ACN) as mobile phases.

The UPLC settings consisted of a sample loading flow rate of 2 μL/min for 8 min followed by a gradient elution with starting with 5% of solvent B and ramping up to 35% over 220 min for total proteomics and 100 min gradient for phosphoproteomics followed by a 10 min wash at 85% B and a 15 min equilibration step at 1% B. The flow rate for the sample run was 300 nL/min with an operating back pressure of about 3800 psi. Full scan survey spectra (m/z 375–1800) were acquired in the Orbitrap with a resolution of 30000 at m/z 400. A data dependent analysis (DDA) was employed in which the five most abundant multiply charged ions present in the survey spectrum were automatically mass-selected, fragmented by collision-induced dissociation (normalized collision energy 35%) and analysed in the LTQ. Dynamic exclusion was enabled with the exclusion list restricted to 500 entries, exclusion duration of 30 s and mass window of 10 ppm.

MASCOT search was used to generate a list of proteins. Peptide identification was by searchers against the SwissProt database (version 2013-2014) restricted to human entries using the Mascot search engine (v 2.5.0, Matrix Science, London, UK). The parameters included trypsin as digestion enzyme with up to two missed cleavages permitted, carbamidomethyl (C) as a fixed modification and Pyro-glu (N-term), Oxidation (M) and Phospho (STY) as variable modifications. Datasets were searched with a mass tolerance of ±5 ppm and a fragment mass tolerance of ±0.8 Da. Label-free quantification was carried out with Pescal as outline above.

### Phosphoproteomic Data Processing and Analysis

Data normalization, log2 transformation, and standardization were performed for downstream analysis. Sample similarity was evaluated using Spearman correlation and principal component analysis (PCA), leading to the exclusion of outliers. Differentially phosphorylated proteins were identified using a t-test, with BH method applied for multiple testing. Proteins with a p-value < 0.05 and an absolute log2 fold change > 1 were considered significantly differentially phosphorylated. Functional enrichment analysis on Gene Ontology (GO)^108^ and Kyoto Encyclopedia of Genes and Genomes (KEGG)^109^ were conducted using the clusterProfiler (v4.10.0)^110^ package in R (v4.3.2). Additionally, kinase enrichment analysis for the differentially phosphorylated proteins was performed using the Kinase Library^111^. Differentially phosphorylated proteins common to the RBD vs WT and WT+Byl vs WT comparisons were identified, and homologous gene conversion was performed using homologene (v1.4.68.19.3.27) package.

Visualization of volcano plots and expression heatmaps was done using ggplot2 (v3.4.4) and heatmap (v1.0.12), respectively. All the phosphoproteomic data processing, statistical analyses, and visualizations were performed in R.

### RNAseq analysis

Gene expression quantification was performed using the Salmon algorithm^112^ to achieve transcript-level quantification. Differential expression analysis was conducted with the edgeR R package^113^, applying its statistical framework to identify significantly differentially expressed genes. Over-representation analysis was carried out using the clusterProfiler suite of R packages^110,114^ to identify enriched pathways and functional categories.

### Collagen gel contraction assays

To assess force-mediated collagen remodeling, 4 × 10⁴ murine CAFs, or 8 × 10⁴ human CAFs (hCAFs), were embedded in 500 µL of Rat tail type I collagen (Corning) solution prepared by mixing collagen stock with 2.5% NaOH (1N) and 16.6 ng TGF-β1 in PBS under sterile conditions at 4°C. Once the gel was set, cells were maintained in DMEM+10%FBS. Gel contraction was monitored daily by taking photographs of the gels. The gel contraction value refers to the contraction observed after 3 days for murine CAFs or 5 days for hCAFs. Contraction index was calculated as the percentage reduction in gel surface area relative to the initial area using ImageJ software.

Where indicated, alpelisib (0.5 μM), Breaker (1.0 μM) or DMSO vehicle was added to the culture medium immediately after collagen gel polymerization and maintained throughout the assay.

### Morphological analysis of CDM fibers

Morphological features were extracted from the microscopy images using a pipeline of traditional image processing steps implemented in Matlab© (The Mathworks™, Natick, USA). The original color images were first converted to grayscale and then processed with a Gaussian low-pass filter (filter size = 8, standard deviation = 3) to reduce noise. The filtered image was subtracted from the grayscale image to remove background noise and subsequently thresholded. The thresholded image was morphologically thinned, ensuring each region was reduced to a width of one pixel, and regions of connected pixels were identified and labeled. Regions with a major axis smaller than 20 pixels were excluded from the analysis. For each image, the following morphological metrics were extracted: number of objects, length, and width of each object. It is important to note that one object does not necessarily correspond to a single fiber. A single fiber may appear as two or more objects if it is broken, or multiple fibers may be merged into a single object if they are intersecting or in close proximity. However, as the average number of objects extracted per image was in the thousands, the extracted metrics can be considered representative of the fibers in the images.

The code used for this analysis is available publicly at the following repository: https://github.com/reyesaldasoro/RBD_Cells. The repository will be made public upon the acceptance of the manuscript.

### CDM stiffness

The elastic modulus of matrices produced by fibroblasts was quantitatively evaluated using Atomic Force Microscopy (AFM). Fibroblasts were cultured on cover glass slides, allowing them to synthesize extracellular matrix (ECM) directly on the coverslips. These prepared samples were then placed into a BIO-AFM system, specifically designed for integration with an inverted optical microscope (Nikon TE2000). The BIO-AFM system utilized a V-shaped silicon nitride probe (nominal spring constant k = 0.03 N/m) with a four-sided pyramidal tip (Bruker AFM Probes) to ensure precise force measurements.

To minimize localized variability and enhance data representativeness, no more than three measurements were taken from a single field of view for mechanical characterization. For each matrix type, stiffness was determined by analyzing data collected from ten different points, with measurements performed in triplicate. Force-displacement (F vs. z) curves were obtained at each point using an amplitude of 10 µm and a frequency of 1 Hz.

The elastic modulus was derived from the force-displacement curves through analysis using the Hertz model, as previously described^115,116^. For each point, five force-displacement curves were recorded, maintaining an indentation depth of 500 nm. Finally, the elastic modulus for each fibroblast-derived ECM subpopulation was calculated. This analysis was based on a minimum of fifty measurements per independent sample, with at least three independent matrices analyzed for each condition (n ≥ 3).

### BMDM isolation and differentiation

Bone marrow-derived macrophages (BMDMs) were generated from C57BL/6 mice. Briefly, bone marrow cells were isolated from the femurs and tibias of the hind limbs by flushing the bones with culture medium under aseptic conditions. The resulting cell suspension was filtered through a 70 µm strainer (Corning), centrifuged and resuspended in complete DMEM supplemented with macrophage colony-stimulating factor (M-CSF; 20 ng/mL; BioLegend) to promote macrophage differentiation. Cells were maintained at 37°C and 5% CO₂ for 7 days. After differentiation, macrophages were fluorescently marked with ViaFluor SE cell prolifera-tion staining (Biotium), collected and seeded onto tumor cell–fibroblast co-culture matrices.

### BMDM co-culture and flow cytometry analysis

Bone marrow-derived macrophages (BMDMs) were generated from mouse bone marrow cells and co-cultured with WT or RBD CAFs for 24 h. BMDMs were collected and analyzed by flow cytometry for macrophage polarization-associated markers, including arginase and iNOS. Cells were stained with the indicated antibodies according to standard flow cytometry procedures, and data were acquired using Aurora 5L Spectral flow cytometer (Cytek Biosciences, Fremont, CA, USA RRID: SCR_019826). Data was acquired using SpectroFlo v3.3.0 software (Cytek Biosciences, Fremont, CA, USA RRID: SCR_025494) and analyzed with Infinicyt v3.0 (BD Biosciences, RRID: SCR_026033). Antibody references and dilutions are listed in Supplementary Table S6.

### Confocal Microscopy

All measurements were performed on a Leica TCS SP8 confocal laser scanning microscope, using a 1.40 NA 63× objective (HCX PL APO CS2 63/1.40 Oil Leica Microsystems). For focal adhesion directionality and colocalization analyses 1024x1024 images were acquired exciting with a WLL (White Light Laser) at 488 and 568 nm for the fibronectin and focal adhesions respectively with a zoom factor of 2 and images were obtained in the axial plane every 0.3 µm focusing on the focal adhesions solely. Directionality analysis was performed at the equatorial plane where lateral focal adhesions were observed in contact with fibrils while Colocalization analysis was performed at the bottom of the cells were focal adhesions and fibrils overlap in the section. Data analysis was performed using Fiji App1 and more specifically using the “Directionality” Plugin for the directionality analysis and JacoP Plug in for the colocalization analysis.

### Fluorescence Lifetime Imaging Microscopy (FLIM) Analysis

To assess ECM accessibility and binding properties, we employed Fluorescence Lifetime Imaging Microscopy (FLIM) using the molecular rotor dye DCVJ (9-(2,2-Dicyanovinyl)julolidine) (Sigma-Aldrich), which exhibits viscosity-dependent fluorescence lifetimes. A concentration gradient of DCVJ (100 nM–1 μM) was applied to decellularized extracellular matrix (CDM) fibers generated by WT and RBD fibroblasts. FLIM images (512 × 512 resolution) were acquired using a Leica TCS SP8 confocal laser scanning microscope equipped with a 63×/1.40 NA oil-immersion objective (HCX PL APO CS2 63/1.40 Oil Leica Microsystems) and SP8 Falcon Module.

### Phasor Analysis

Phasor transformations were performed to map fluorescence decay data from each pixel into a graphical representation, facilitating the analysis of complex, multi-exponential lifetime distributions. The phasor coordinates for each pixel were calculated as follows:

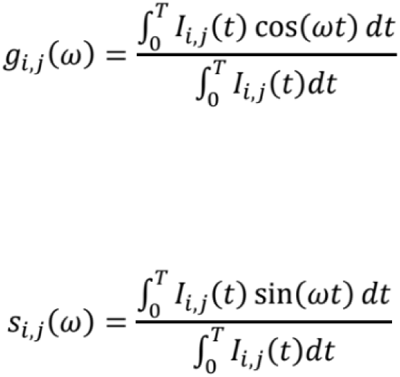

where ω=2πf and f=1/T is the laser repetition rate. The resulting phasors were plotted within the "universal circle," which defines single-exponential decays based on their lifetimes, ranging from τ = 0 at (1,0) to τ = ∞ at (0,0). Mixtures of two components produce phasor points that lie along a straight line connecting the single-component lifetimes, with the position determined by their relative fractional contributions.

Quantitative analysis was performed by defining two single-exponential lifetimes, τ1 = 1.9 ns and τ2 = 0.3 ns, based on control measurements in glycerol. A linear combination model was applied to calculate the fraction of bound versus unbound molecules within ECM fibers using the relative distances from each phasor point to τ1 and τ2. Lineweaver-Burk plots were generated to determine binding constants and relative binding site availability between WT and RBD CDMs.

### Western blot analysis

For Western blot analysis, cells were scraped and lysed in ice-cold RIPA buffer (50 mM Tris-HCl, pH 7.4, 150 mM NaCl, 1% NP-40, 0.5% sodium deoxycholate, 0.1% SDS) supplemented with cOmplete™ mini protease inhibitor cocktail (Roche). Protein concentrations were determined using the Bradford assay according to the manufacturer’s instructions (Biorad). Protein lysates were resolved on 6-18% SDS-PAGE gels (Invitrogen), transferred onto PVDF membranes and immunoblotted with the indicated primary antibodies (Supplementary Table S6) diluted in 5% bovine serum albumin (BSA; NZYtech) in TBS-Tween (0.1% v/v) overnight at 4°C. Primary antibodies were detected using peroxidase-conjugated secondary antibodies (Supplementary Table S6), incubated for 1 h at RT in 5% (w/v) non-fat milk in TBS-Tween (0.1% v/v) for 1 hour at room temperature. The protein bands were visualized using an ECL substrate (Cytiva) on an iBright^TM^ FL1500 (Invitrogen) imaging system.

### RNA isolation, Retrotranscription (RT) and quantitative PCR (qPCR)

Total RNA was extracted using the NZY Total RNA Isolation Kit (NZYtech) following the manufacturer’s protocol. RNA concentration and quality were assessed with a Nanodrop spectrophotometer. 600 ng of RNA were reverse transcribed into cDNA using the PrimeScript™ RT Reagent Kit (Takara) according to the manufacturer’s instructions.

qPCR was performed in triplicate using a QuantStudio™ 5 (384-well block, Thermo Scientific) and PowerTrack SYBR Green Master Mix (Thermo Scientific). Gene expression was normalized to 18S, and relative expression levels were calculated using the 2^−ΔΔCt^ method. Primer sequences are listed in Supplementary Table S7

For gene expression analysis of tumor cells reseeded in decellularized CDMs, 2 × 10⁵ cancer cells were seeded onto CDMs in 6-well plates. RNA was isolated after 6 hours (Snail1, Zeb, Twist) or 24 hours (Vim, N-cadh, E-cadh) as described previously. Prior to RNA isolation, lysates were homogenized using QIAshredder columns (Qiagen).

Gene expression changes were quantified by qPCR and expressed as fold change (FC) relative to the control condition. Mean FC values from three independent biological replicates were log2-transformed and used for heatmap visualization. Data were normalized per gene using row-wise z-score normalization to highlight relative expression patterns across conditions. Heatmaps were generated using Morpheus (Broad Institute), and hierarchical clustering of genes was performed using Pearson correlation and average linkage.

### Proliferation assays in decellularized CDMs

5 × 10^4^ cancer cells were seeded in decellularized CDMs per well in a 24-well plate, at least in triplicates. Proliferation/viability was then assessed at 0, 24 and 48 h using CellTiter-Blue cell viability reagent (Promega). At each timepoint, cells were incubated for 2 h with 50 µl CellTiter-Blue solution (added to tissue culture medium), and then fluorescence was measured in a spectrophotometer using 560/590 (excitation/emission) filter settings.

### Immunofluorescence

Cells were fixed in 4% paraformaldehyde (Sigma)/PBS (v/v) for 15 min at RT, and then permeabilized with 0.1% Triton x100 (Sigma-Aldrich)/PBS (v/v) for 5 min at RT, blocked with 3% BSA (w/v) (NZYTech) for 1 h at RT and stained with primary antibodies overnight in humidified chambers at 4°C. Sections were then washed 3 times with PBS for 5 min, incubated with secondary antibodies and phalloidin, depending on the staining, for 1 h at RT, washed 3 times in PBS for 5 min and mounted using ProLong Gold Antifade reagent with DAPI (Invitrogen). Antibody references and dilutions are listed in Supplementary Table S6. For IF of cancer cells reseeded in decellularized CDMs, 1.2 × 10⁵ cancer cells were seeded onto CDMs in 6-well plates. After 24 hours, cells were fixed with 4% PFA at room temperature for 20 minutes. The IF protocol described earlier was followed, except for permeabilization, which was performed with 0.5% Triton X-100 for 10 minutes.

### Histology and Analysis

Mice were sacrificed and lung tissues were immediately removed and fixed overnight in 4% PFA. Lungs were transferred to 70% ethanol and processed for paraffin embedding. Tissue sections were cut at 4 µm and stained with H&E or immunostained with the indicated antibodies (Supplementary Table S6). *Tumor burden quantification*: Lung and tumor areas were quantified on H&E-stained sections. Images of each lung lobe were acquired using an Olympus Bx51 microscope equipped with a DE74 camera and a 1× objective. Lung and tumor areas were measured using ImageJ.

#### Phospho-histone H3 (PHH3) quantification

Cancer cell nuclei and PHH3-positive nuclei were counted using the Fiji multi-point tool. The proportion of PHH3-positive cells was calculated by dividing the number of PHH3-positive nuclei by the total number of cancer cells in each tumor cluster.

#### α-SMA, fibronectin, Alcian Blue and Sirius Red quantification

Positive staining area was quantified using intensity threshold-based segmentation and normalized as indicated for each analysis. Cancer cells were counted manually using the Fiji multi-point tool.

### Statistical Analysis

Data are presented as mean ± SEM unless otherwise indicated. Statistical analyses were performed using GraphPad Prism 10. For comparisons between two groups, statistical significance was determined using unpaired two-tailed Student’s t-test unless otherwise stated. Survival curves were analyzed using the log-rank test. The number of biological replicates, animals or independent experiments is indicated in the corresponding figure legends. Statistical significance was defined as p < 0.05.

## SUPPLEMENTARY FIGURES

**Supplementary Figure 1.**
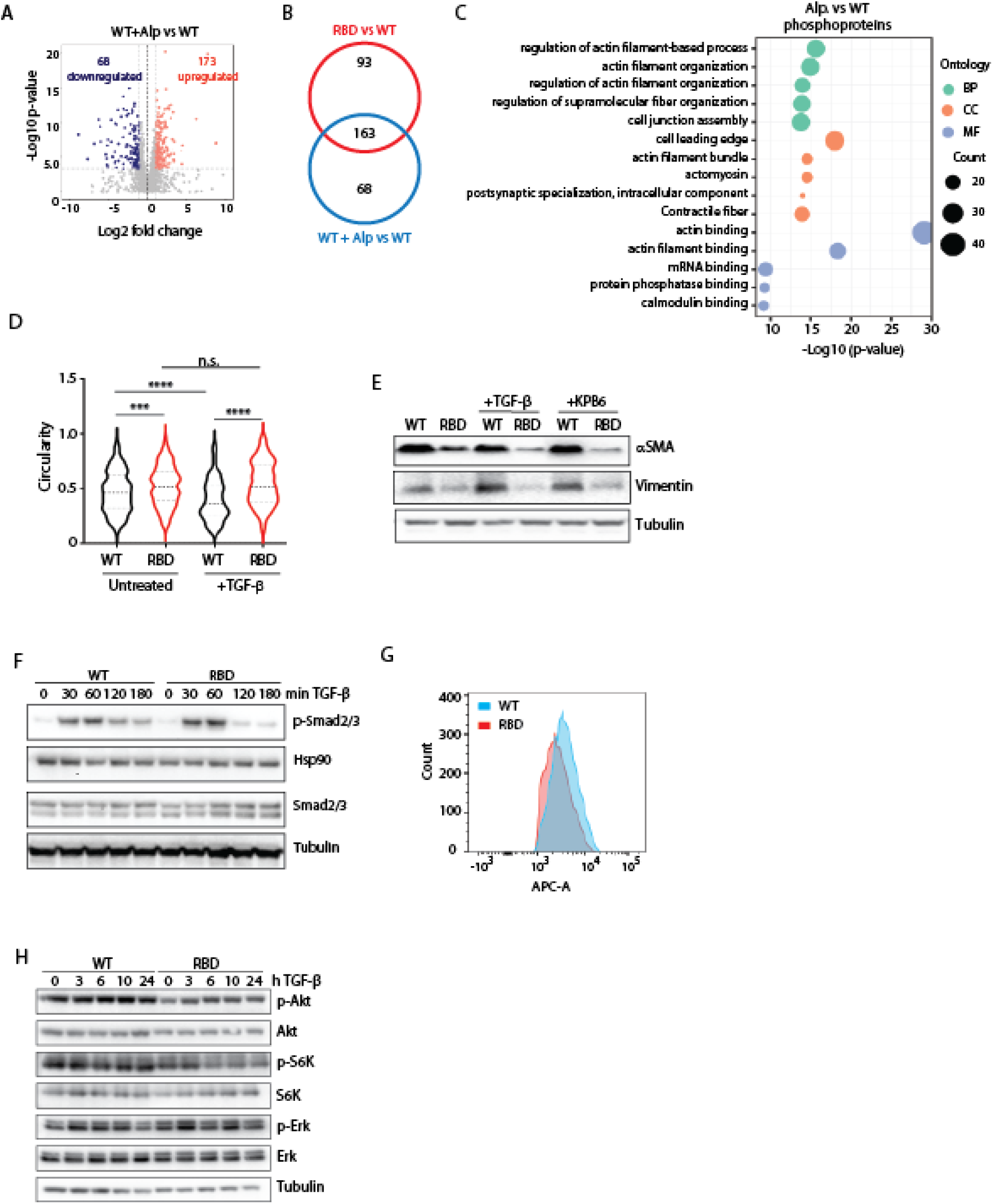
Pharmacological and signaling validation of RAS–PI3K-dependent CAF activation defects. (A) Volcano plot showing differentially phosphorylated peptides identified by phosphoproteomic analysis of lung tumors from alpelisib-treated WT mice compared with vehicle-treated WT controls. Significantly downregulated and upregulated phosphopeptides are shown in blue and red, respectively. (B) Venn diagram showing the overlap between proteins differentially phosphorylated in *Pik3ca*^RBD/−^ *versus Pik3ca*^WT/−^ tumors and in alpelisib-treated WT tumors versus WT controls, indicating a shared RAS–PI3K-dependent signaling response *in vivo*. (C) Gene Ontology enrichment analysis of significantly altered phosphopeptides in alpelisib-treated WT tumors versus controls, highlighting pathways related to actin filament organization, cell adhesion, cytoskeletal regulation and phosphatase activity. Dot size indicates the number of proteins associated with each term; color denotes ontology category. (D) Quantification of cellular circularity in WT and RBD fibroblasts under basal conditions or following TGF-β stimulation, showing increased circularity in RAS–PI3K-deficient fibroblasts after activation. (E) Immunoblot analysis of α-SMA and vimentin protein levels in WT and RBD fibroblasts following stimulation with TGF-β or KPB6-CM. α-Tubulin was used as loading control. (F) Immunoblot analysis of SMAD2/3 phosphorylation and total SMAD2/3 levels in WT and RBD fibroblasts following TGF-β stimulation for the indicated times. α-Tubulin was used as loading control. (G) Flow cytometry analysis of surface TGFβRII receptor expression in WT and RBD fibroblasts. (H) Immunoblot analysis of AKT, S6K and ERK phosphorylation, together with total AKT, S6K and ERK levels, in WT and RBD fibroblasts following TGF-β stimulation for the indicated times. α-Tubulin was used as loading control. Data are shown as violin plots with median and quartiles where applicable. Statistical significance was determined using Student’s t-test unless otherwise indicated. ***p < 0.001; ****p < 0.0001; n.s., not significant.

**Supplementary Figure 2.**
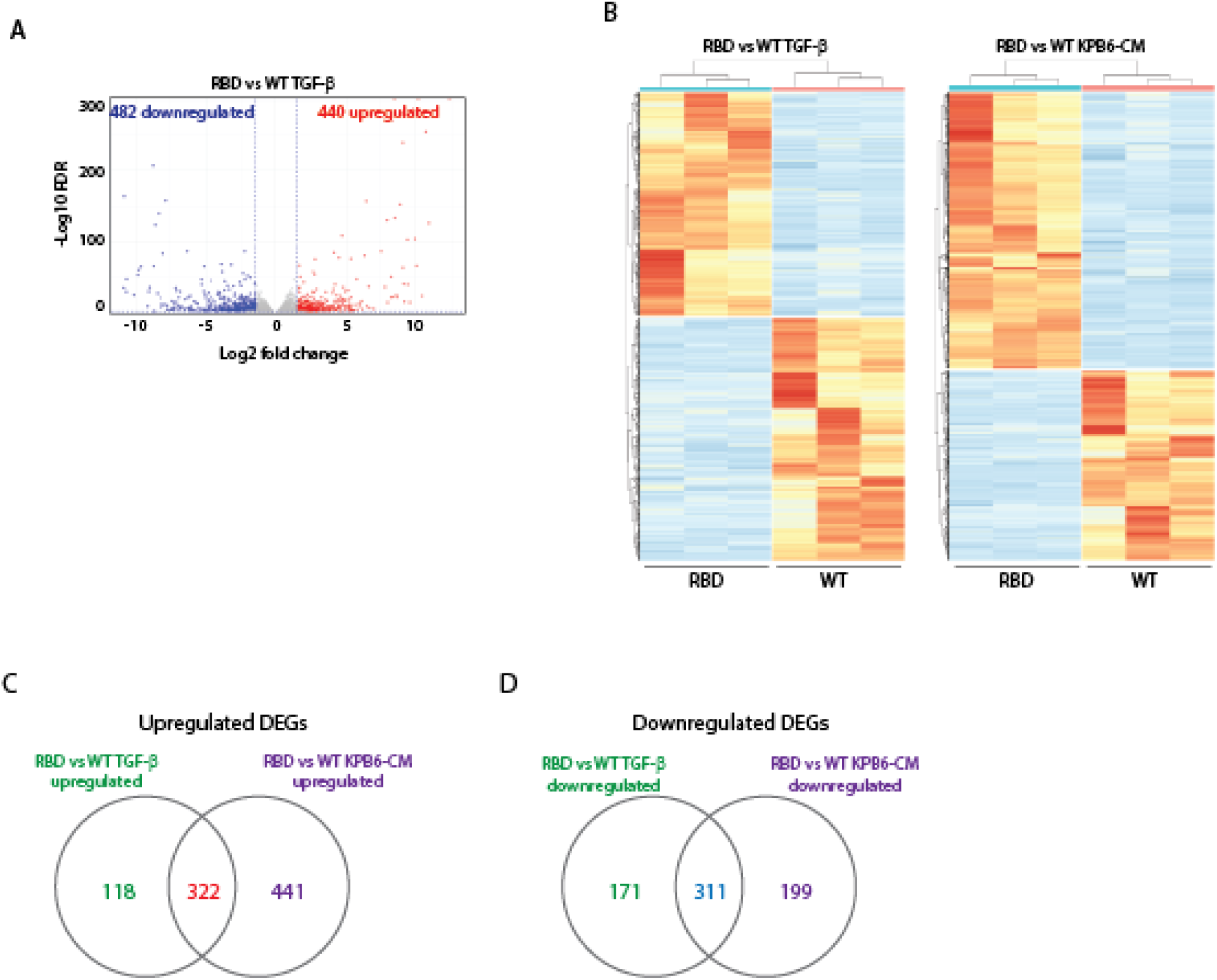
RAS–PI3K disruption imposes a conserved transcriptional program across profibrotic and tumor-derived CAF activation cues. (A) Volcano plot showing differentially expressed genes in RBD versus WT CAFs following TGF-β stimulation. Significantly downregulated and upregulated genes are shown in blue and red, respectively. (B) Heatmaps showing differentially expressed genes in RBD versus WT CAFs following stimulation with TGF-β or KPB6-CM. Each column represents an independent biological replicate. (C) Venn diagram showing the overlap between genes upregulated in RBD versus WT CAFs following TGF-β stimulation and genes upregulated in RBD versus WT CAFs following KPB6-CM stimulation. (D) Venn diagram showing the overlap between genes downregulated in RBD versus WT CAFs following TGF-β stimulation and genes downregulated in RBD versus WT CAFs following KPB6-CM stimulation. Differentially expressed genes were identified from RNA-seq analysis of WT and RBD CAFs stimulated with either TGF-β or KPB6-CM. Heatmaps show normalized expression values. The overlap between transcriptional responses indicates that loss of RAS–PI3K signaling imposes a shared directional bias across distinct CAF-activating stimuli.

**Supplementary Figure 3.**
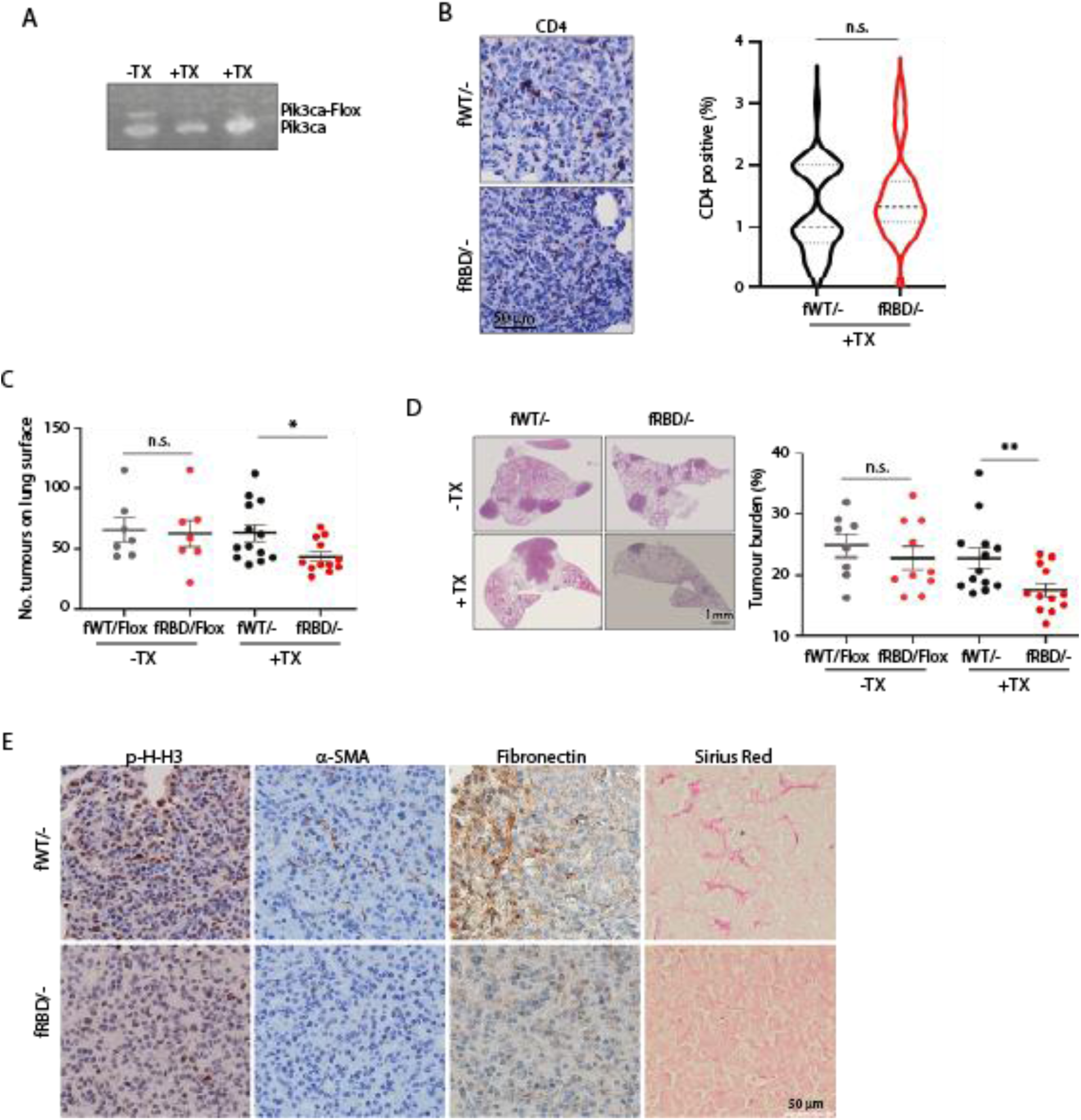
CAF-specific RAS–PI3K disruption reduces tumor growth and stromal remodeling in autochthonous KRAS-driven lung tumors. (A) PCR analysis confirming recombination of the floxed *Pik3ca* allele in lung fibroblasts isolated from *Pik3ca*^fRBD/Flox^ mice following tamoxifen treatment. (B) Representative CD4 immunohistochemistry and quantification of CD4-positive T cells in tumors from tamoxifen-treated *Pik3ca*^fWT/−^ and *Pik3ca*^fRBD/−^ mice following intravenous KPB6 cell injection. Scale bar, 50 μm. (C) Quantification of the number of tumors on the lung surface in *Pik3ca*^fWT/−^ and *Pik3ca*^fRBD/−^ mice crossed with the KRAS^LA2^ model, in the absence or presence of tamoxifen-induced recombination. (D) Representative H&E-stained lung sections and quantification of tumor burden in *Pik3ca*^fWT/−^;KRAS^LA2^ and *Pik3ca*^fRBD/−^;KRAS^LA2^ mice in the absence or presence of tamoxifen-induced recombination. Tumor burden is expressed as tumor area relative to total lung area. Scale bar, 1 mm. (E) Representative immunohistochemistry and histological staining of tumors from tamoxifen-treated *Pik3ca*^fWT/−^;KRAS^LA2^ and *Pik3ca*^fRBD/−^;KRAS^LA2^ mice, showing phospho-histone H3, α-SMA, fibronectin and Sirius Red staining. Scale bar, 50 μm. Data are shown as mean ± SEM or violin plots with median and quartiles, as indicated. Statistical significance was determined using Student’s t-test unless otherwise indicated. p < 0.05; *p < 0.01; n.s., not significant.

**Supplementary Figure 4.**
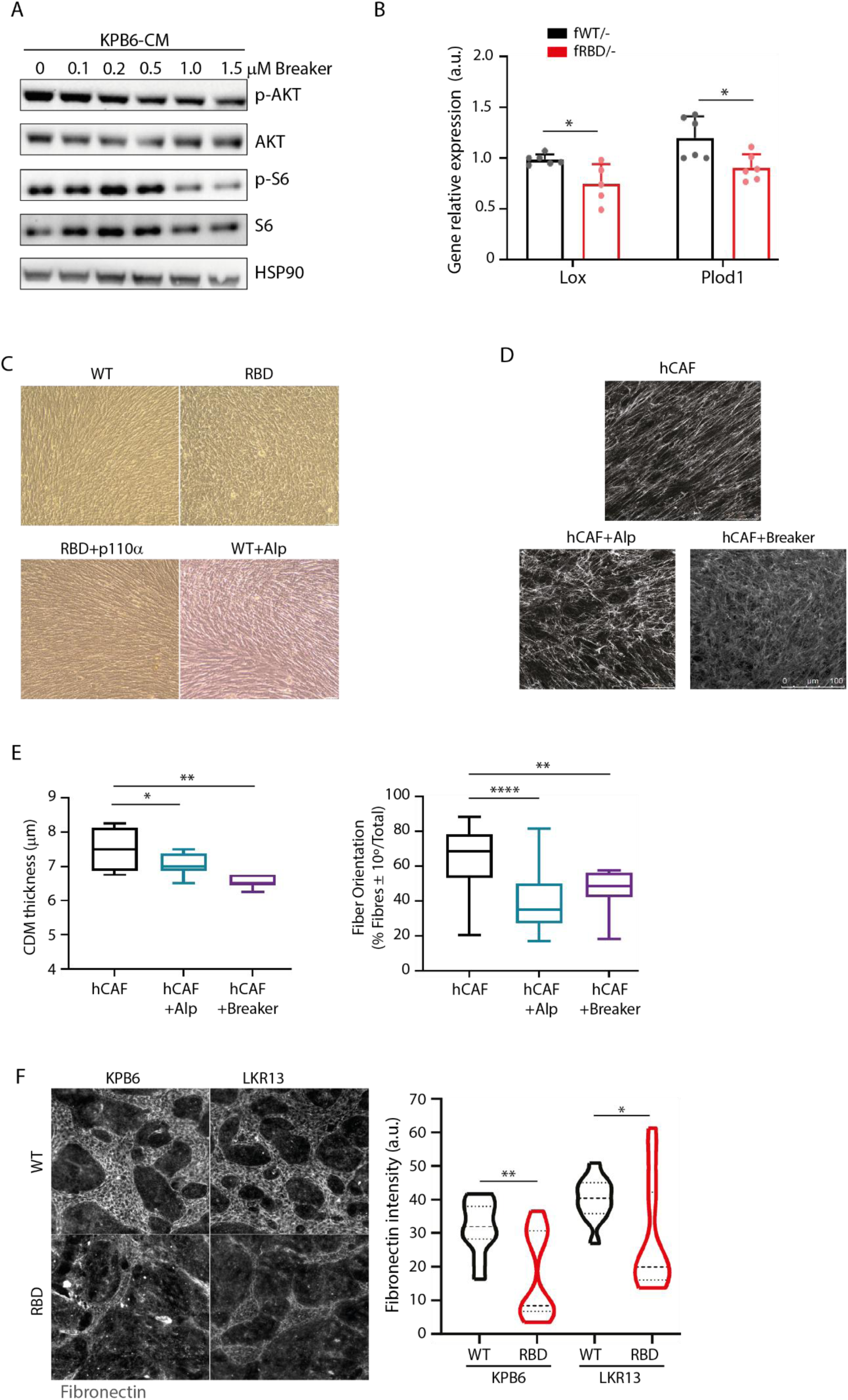
Genetic and pharmacological disruption of RAS–PI3K signaling impairs CAF contractility and matrix organization across murine and human systems. (A) Immunoblot analysis of AKT and S6 phosphorylation in CAFs stimulated with KPB6-CM and treated with increasing concentrations of the RAS–PI3K interaction Breaker. Total AKT, total S6 and HSP90 were used as controls. (B) Quantitative PCR analysis of Lox and Plod1 expression in whole tumors from tamoxifen-treated *Pik3ca*^fWT/−^ and *Pik3ca*^fRBD/−^ mice, showing reduced expression of collagen maturation enzymes upon CAF-specific RAS–PI3K disruption. (C) Representative transmitted-light images of WT and RBD fibroblast cultures during CAF-derived matrix generation, showing comparable cell persistence throughout the matrix-production period. (D) Representative fibronectin immunofluorescence images of CAF-derived matrices generated by human lung cancer-associated fibroblasts treated with vehicle, alpelisib or Breaker. (E) Quantification of matrix thickness and fiber alignment in human CAF-derived matrices treated with vehicle, alpelisib or Breaker. (F) Representative fibronectin immunofluorescence images and quantification of fibronectin organization in matrices generated by WT or RBD fibroblasts co-cultured with two independent KRAS-mutant lung cancer cell lines, KPB6 and LKR13. RBD fibroblast-derived matrices displayed reduced fibronectin fibrillar organization compared with WT controls under both tumor co-culture conditions. Data are shown as bar graphs, box-and-whisker plots or violin plots as indicated. Statistical significance was determined using Student’s t-test unless otherwise indicated. p < 0.05; *p < 0.01; n.s., not significant.

**Supplementary Figure 5.**
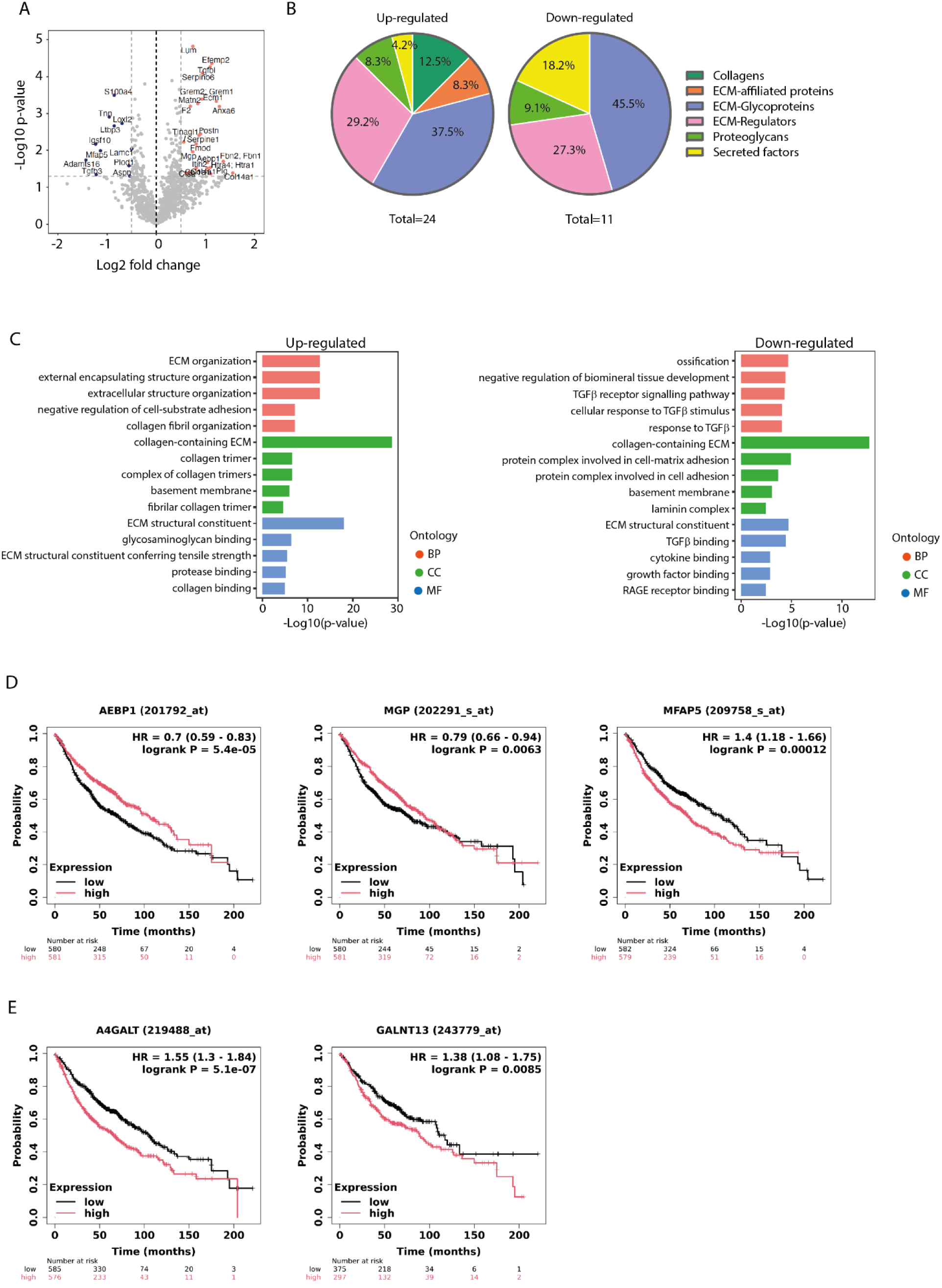
RAS–PI3K-dependent matrisome remodeling reveals glycoprotein and glycosylation-associated alterations linked to lung cancer outcome. (A) Volcano plot showing differentially expressed proteins identified by quantitative proteomic analysis of decellularized CAF-derived matrices generated by RBD *versus* WT fibroblasts. Significantly deregulated matrisome-associated proteins are highlighted and annotated. (B) Pie charts showing the distribution of matrisome categories among proteins upregulated or downregulated in RBD-derived matrices compared with WT-derived matrices. ECM glycoproteins are preferentially represented among downregulated components, whereas collagens and ECM-affiliated proteins are more prominently represented among upregulated components. (C) Gene Ontology enrichment analysis of significantly upregulated and downregulated matrisome-associated proteins in RBD versus WT CAF-derived matrices. Enriched terms highlight structural collagen-associated pathways among upregulated proteins and cell–matrix adhesion, TGF-β signaling and protein interaction networks among downregulated proteins. (D) Kaplan–Meier survival analysis of lung adenocarcinoma patients stratified according to expression of selected RAS–PI3K-regulated ECM glycoprotein-associated genes identified in the CAF-derived matrisome analysis. Hazard ratios and log-rank p values are indicated in each plot. (E) Kaplan–Meier survival analysis of lung adenocarcinoma patients stratified according to expression of selected RAS–PI3K-regulated glycosyltransferase genes. Hazard ratios and log-rank p values are indicated in each plot. For survival analyses, patients were stratified according to high or low gene expression. Statistical significance for proteomic comparisons and enrichment analyses was determined as described in Methods.

**Supplementary Figure 6.**
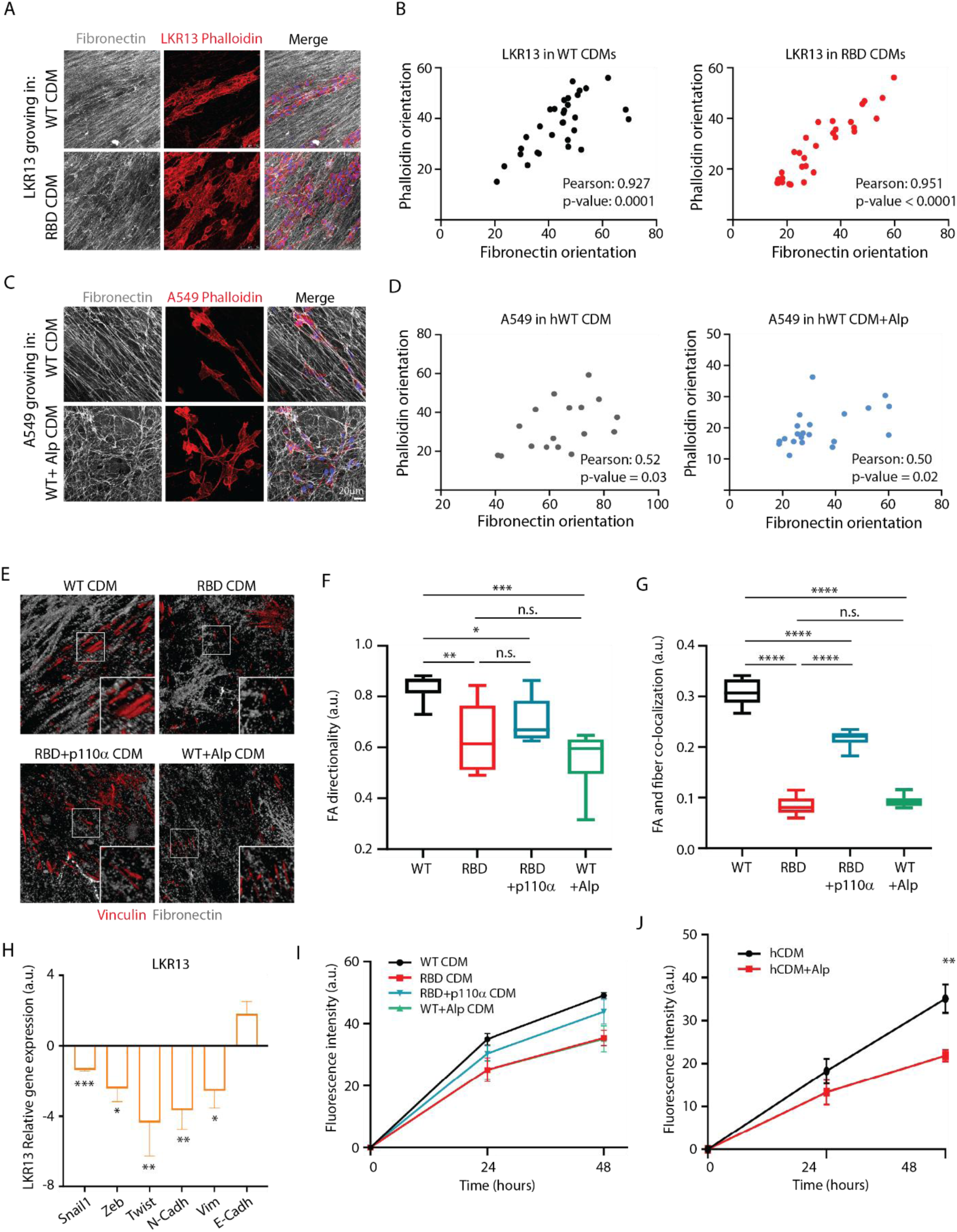
RAS–PI3K-dependent matrix organization controls tumor cell alignment, adhesion and growth across lung cancer models. (A) Representative immunofluorescence images of LKR13 lung cancer cells seeded onto decellularized CAF-derived matrices generated by WT or RBD fibroblasts. Fibronectin fibers are shown in grey and the LKR13 actin cytoskeleton by phalloidin staining in red. Merged images show tumor cell alignment relative to the underlying fibronectin network. (B) Correlation analysis between fibronectin fiber orientation and LKR13 phalloidin orientation in WT-and RBD-derived matrices. Pearson correlation coefficients and p values are indicated. (C) Representative immunofluorescence images of A549 human lung cancer cells seeded onto decellularized matrices generated by human CAFs treated with vehicle or alpelisib. Fibronectin fibers are shown in grey and A549 actin cytoskeleton by phalloidin staining in red. Scale bar, 20 μm. (D) Correlation analysis between fibronectin fiber orientation and A549 phalloidin orientation in control and alpelisib-treated human CAF-derived matrices. Pearson correlation coefficients and p values are indicated. (E) Representative immunofluorescence images of focal adhesions in tumor cells seeded onto CAF-derived matrices generated by WT, RBD or RBD+p110α fibroblasts, or by alpelisib-treated WT fibroblasts. Vinculin-positive focal adhesions are shown in red and fibronectin fibers in grey. Insets show magnified regions of focal adhesion–fibronectin coupling. (F–G) Quantification of focal adhesion directionality (F) and focal adhesion–fibronectin colocalization (G) in the conditions shown in (E). (H) Quantitative PCR analysis of EMT-associated transcription factors and epithelial/mesenchymal markers in LKR13 cells cultured on RBD-derived matrices relative to WT-derived matrices. (I) Proliferation analysis of LKR13 cells cultured on matrices generated by WT, RBD, RBD+p110α or alpelisib-treated WT fibroblasts. (J) Proliferation analysis of A549 cells cultured on human CAF-derived matrices generated in the absence or presence of alpelisib. Data are shown as box-and-whisker plots, bar graphs or mean ± SEM as indicated. Statistical significance was determined using Student’s t-test unless otherwise indicated. *p < 0.05; **p < 0.01; ***p < 0.001; ****p < 0.0001; n.s., not significant.

**Supplementary Figure 7.**
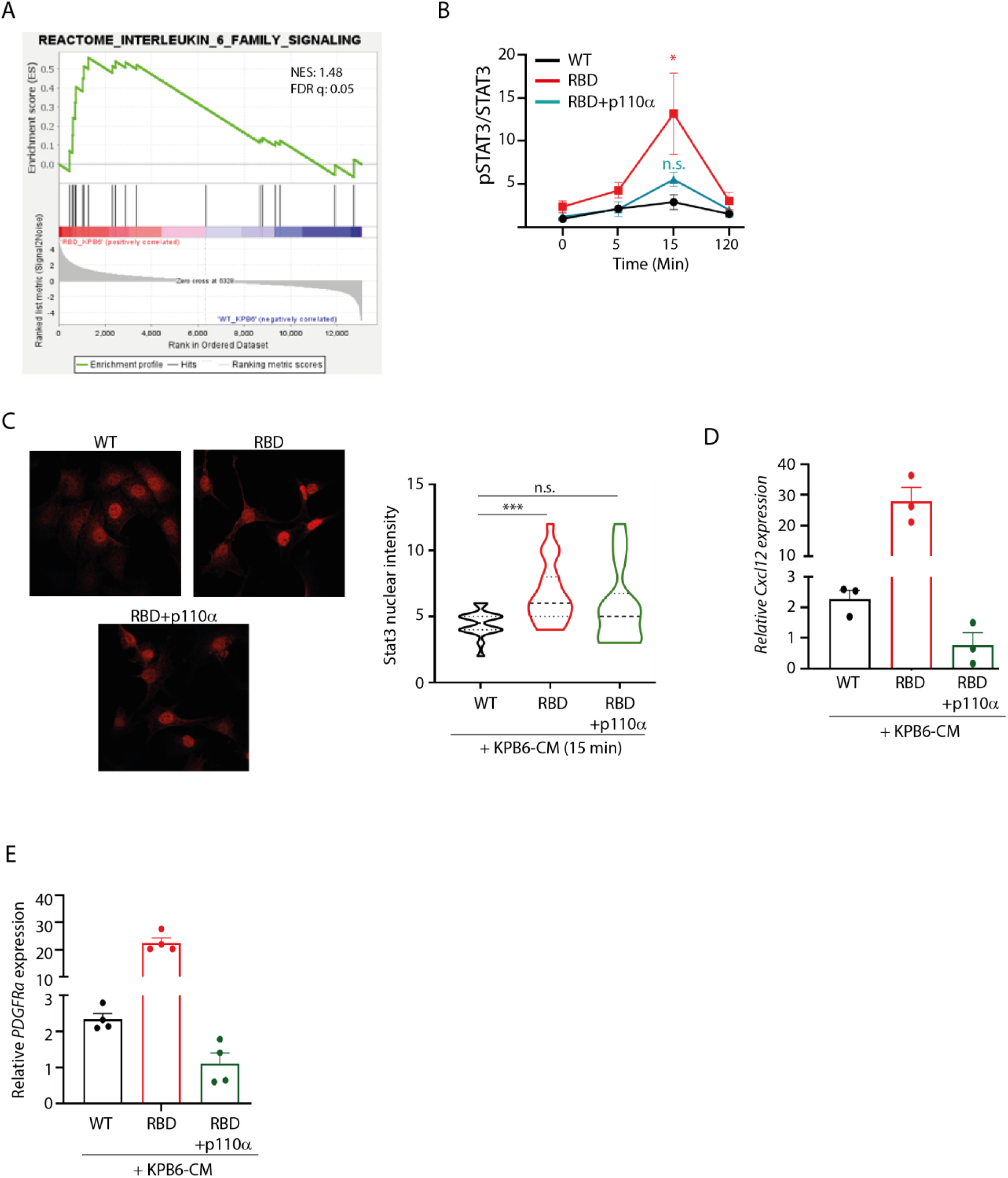
RAS–PI3K disruption promotes sustained STAT3 activation and secretory-state remodeling in CAFs. (A) Gene Set Enrichment Analysis showing enrichment of the REACTOME_INTERLEUKIN_6_FAMILY_SIGNALING pathway in RBD CAFs compared with WT CAFs following stimulation with KPB6-CM. Normalized enrichment score (NES) and false discovery rate (FDR q value) are indicated. (B) Quantification of STAT3 phosphorylation from the immunoblot shown in Figure 7D. p-STAT3 levels were normalized as indicated in the graph and plotted across the indicated time points in WT, RBD and RBD+p110α fibroblasts stimulated with KPB6-CM. (C) Representative immunofluorescence images and quantification of nuclear STAT3 accumulation in WT, RBD and RBD+p110α fibroblasts after 15 min of stimulation with KPB6-CM, showing increased early STAT3 nuclear localization in RBD fibroblasts and reduction upon p110α re-expression. (D) Quantitative PCR analysis of *Cxcl12* expression in WT, RBD and RBD+p110α fibroblasts stimulated with KPB6-CM. (E) Quantitative PCR analysis of *Pdgfrα* expression in WT, RBD and RBD+p110α fibroblasts stimulated with KPB6-CM. Data are shown as mean ± SEM or violin plots with median and quartiles, as indicated. Statistical significance was determined using Student’s t-test unless otherwise indicated. n.s., not significant; *p < 0.05; **p < 0.01; ***p < 0.001; ****p < 0.0001.

