## Supplementary figures and images for "RAS–PI3K signaling promotes myofibroblastic CAF identity and restrains an immunomodulatory stromal program"

### Supplementary Figure 1S

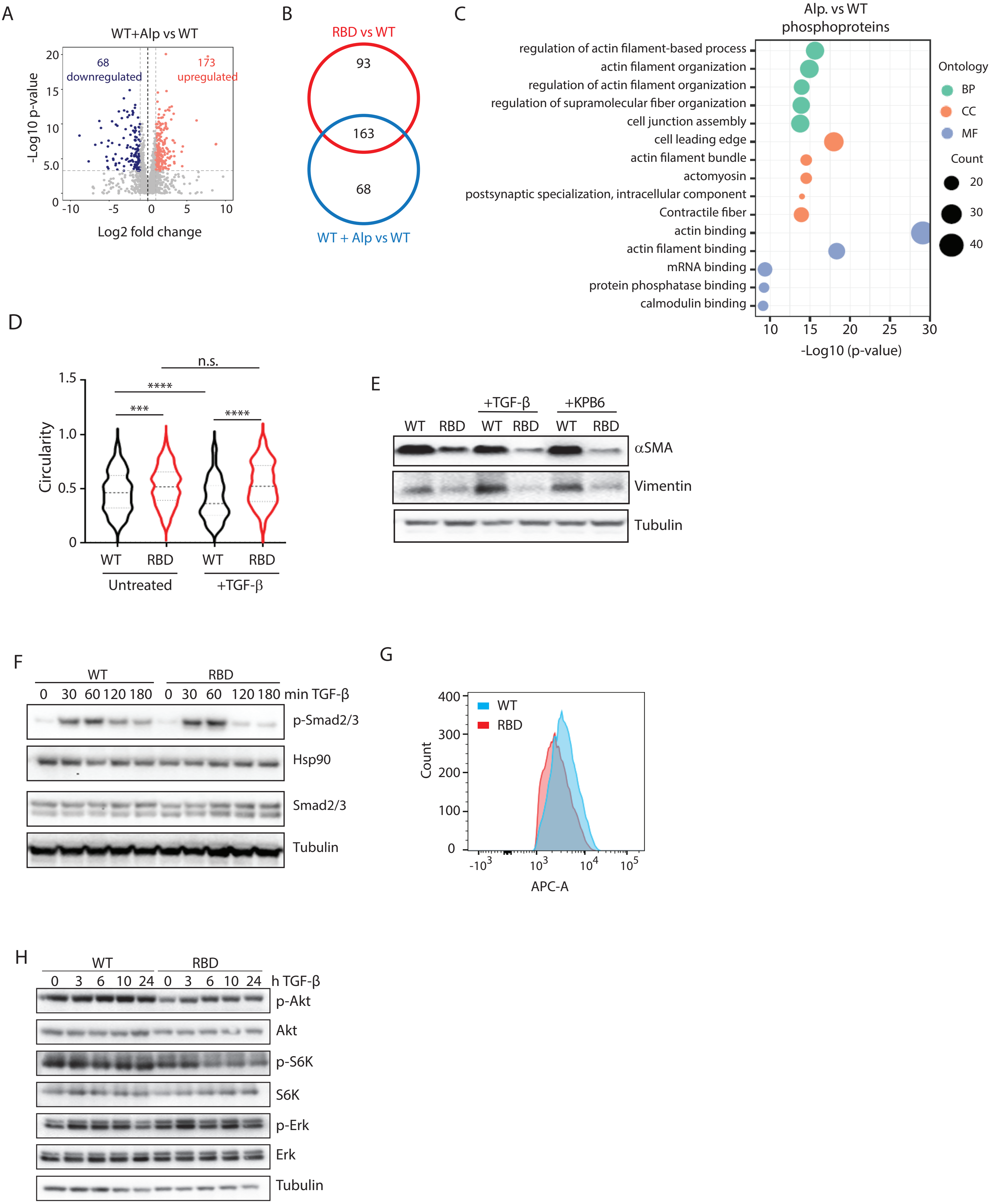

### Supplementary Figure 2S

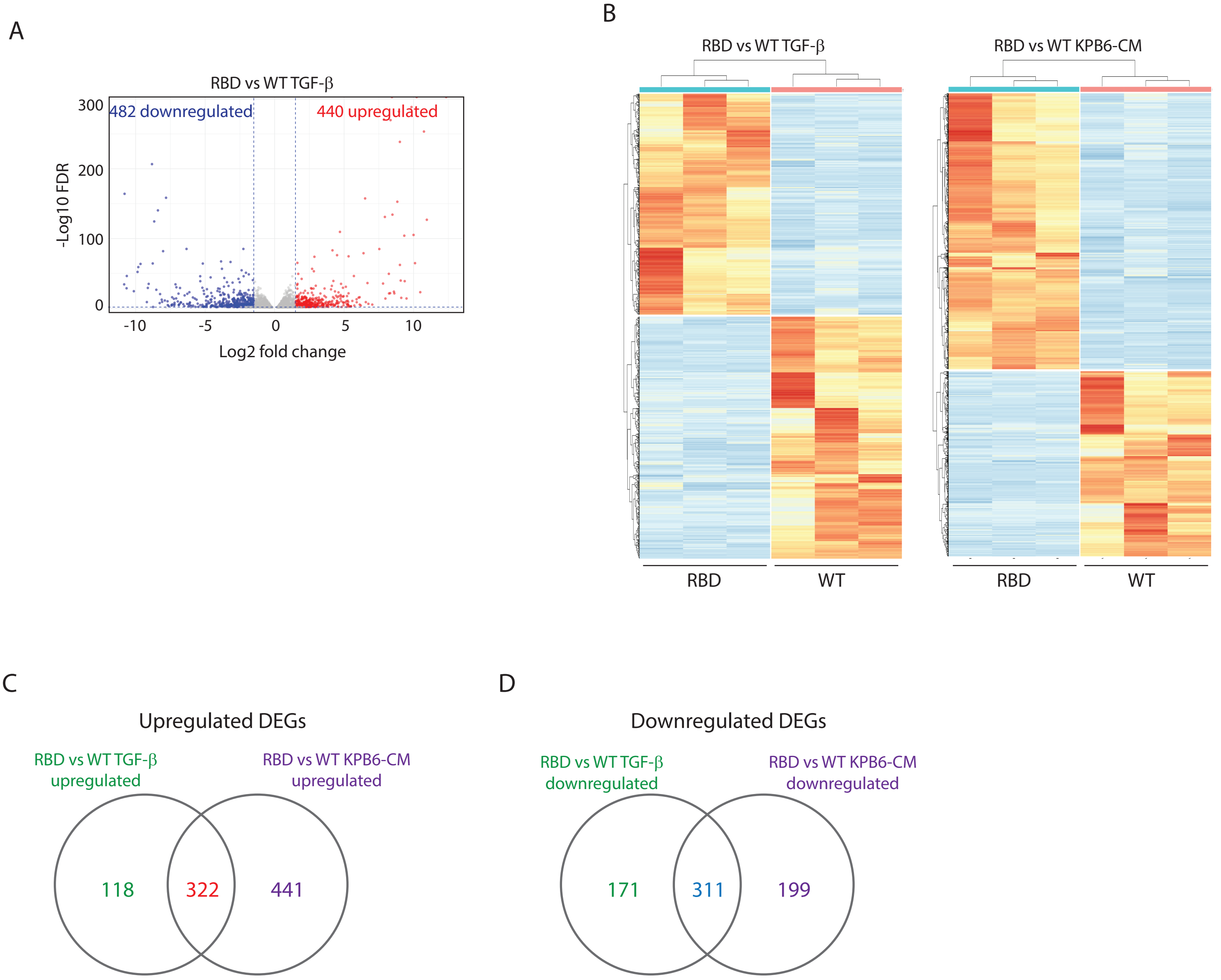

### Supplementary Figure 3S

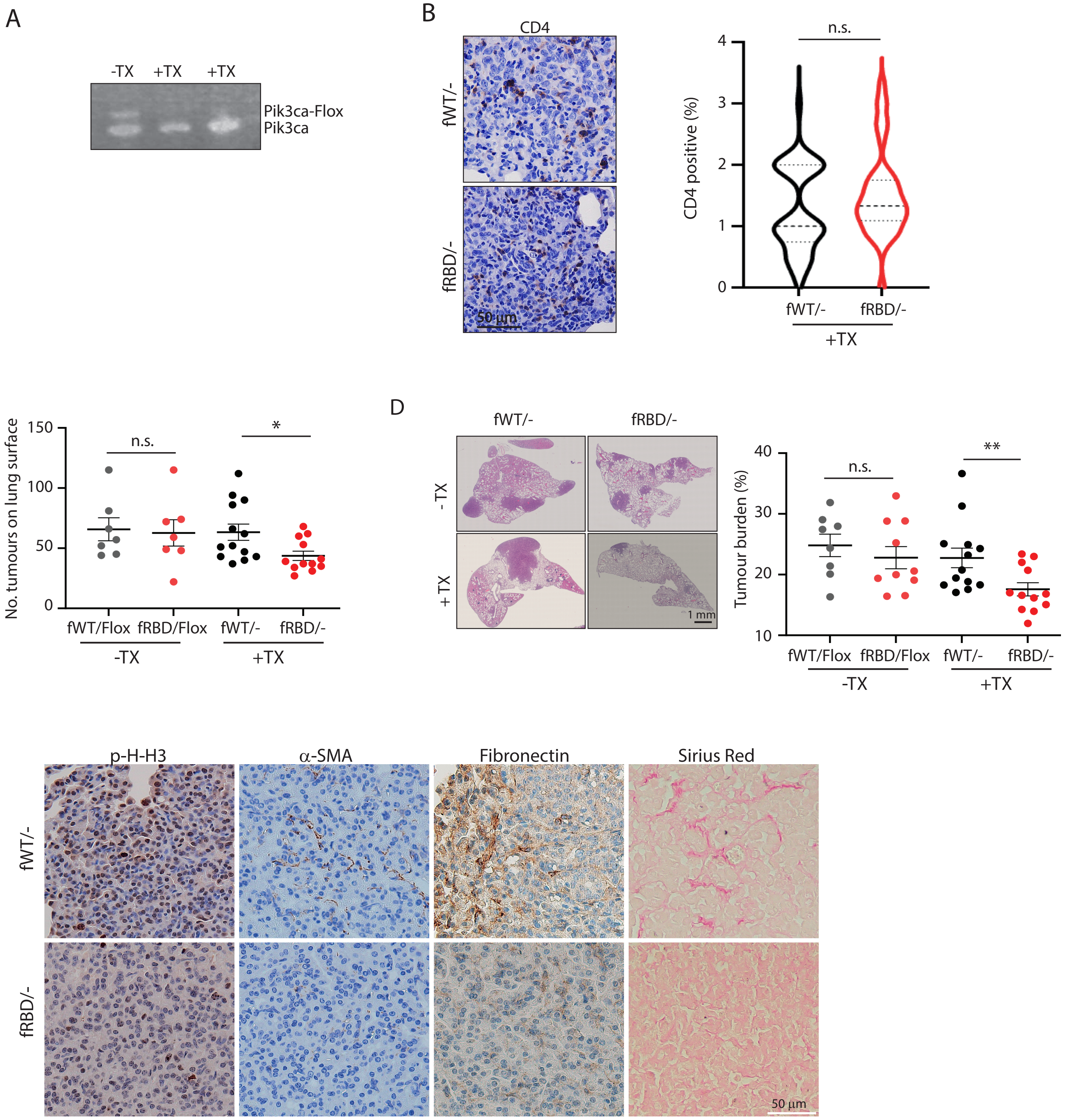

### Supplementary Figure 4S

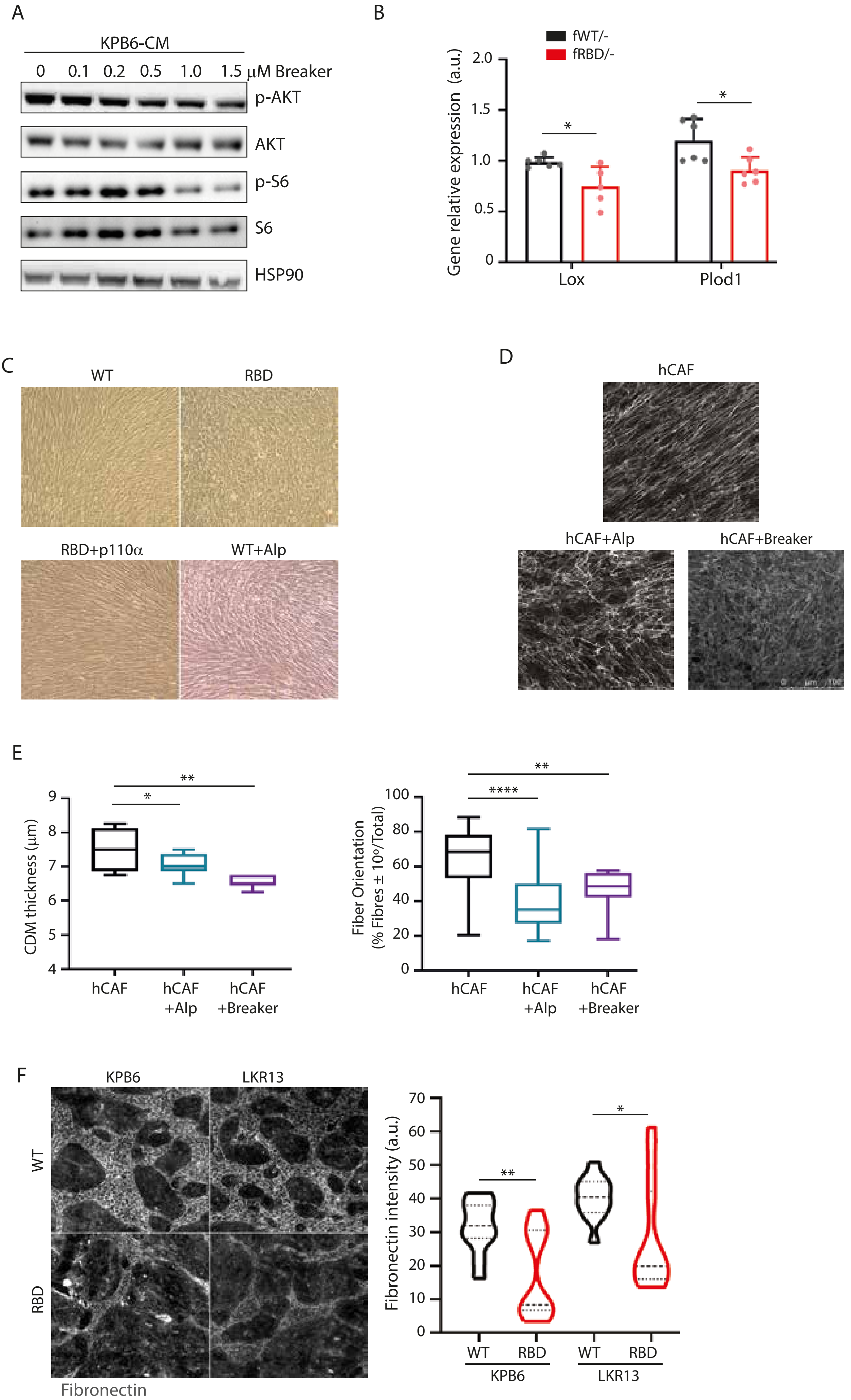

### Supplementary Figure 5S

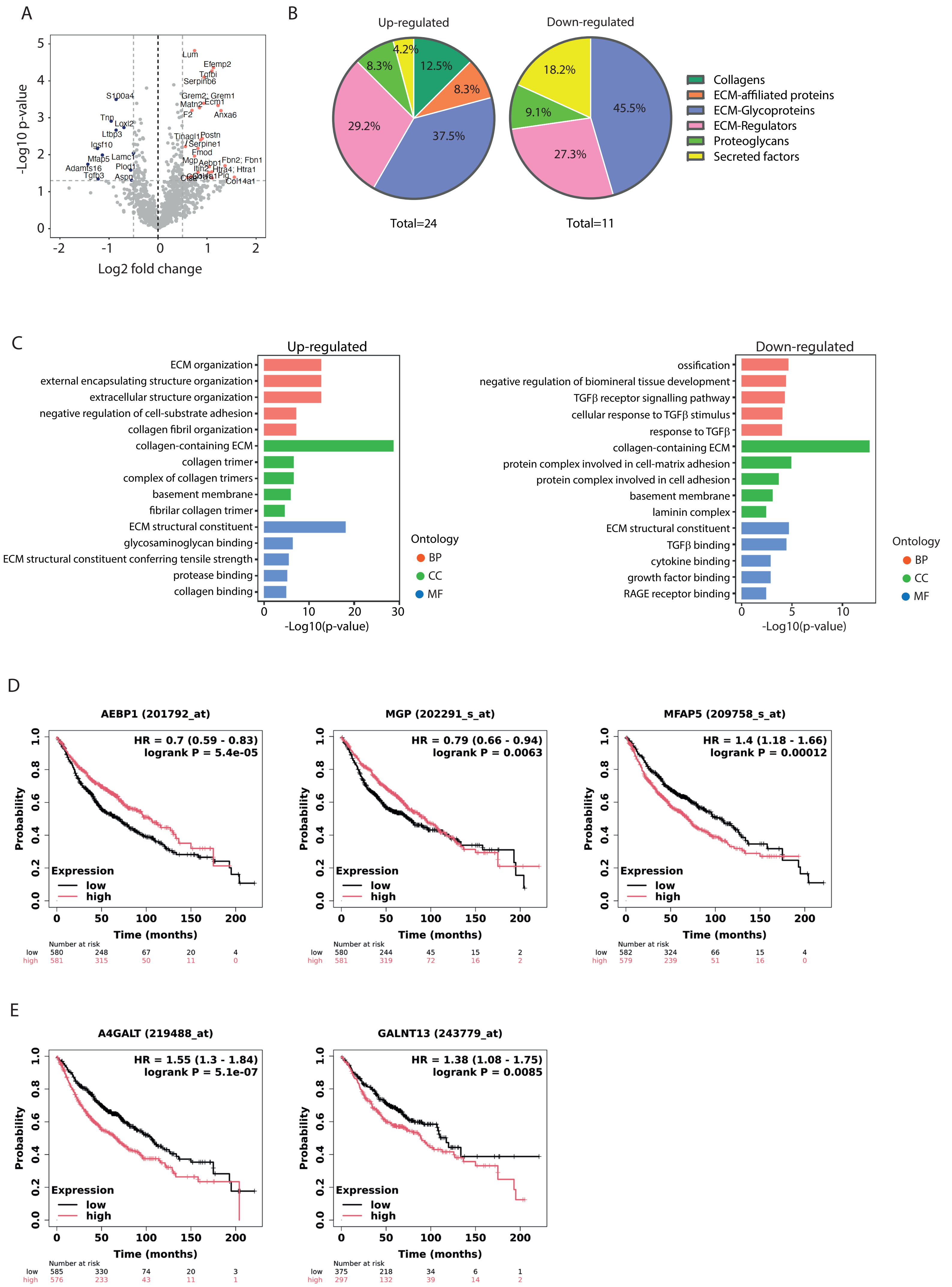

### Supplementary Figure 6S

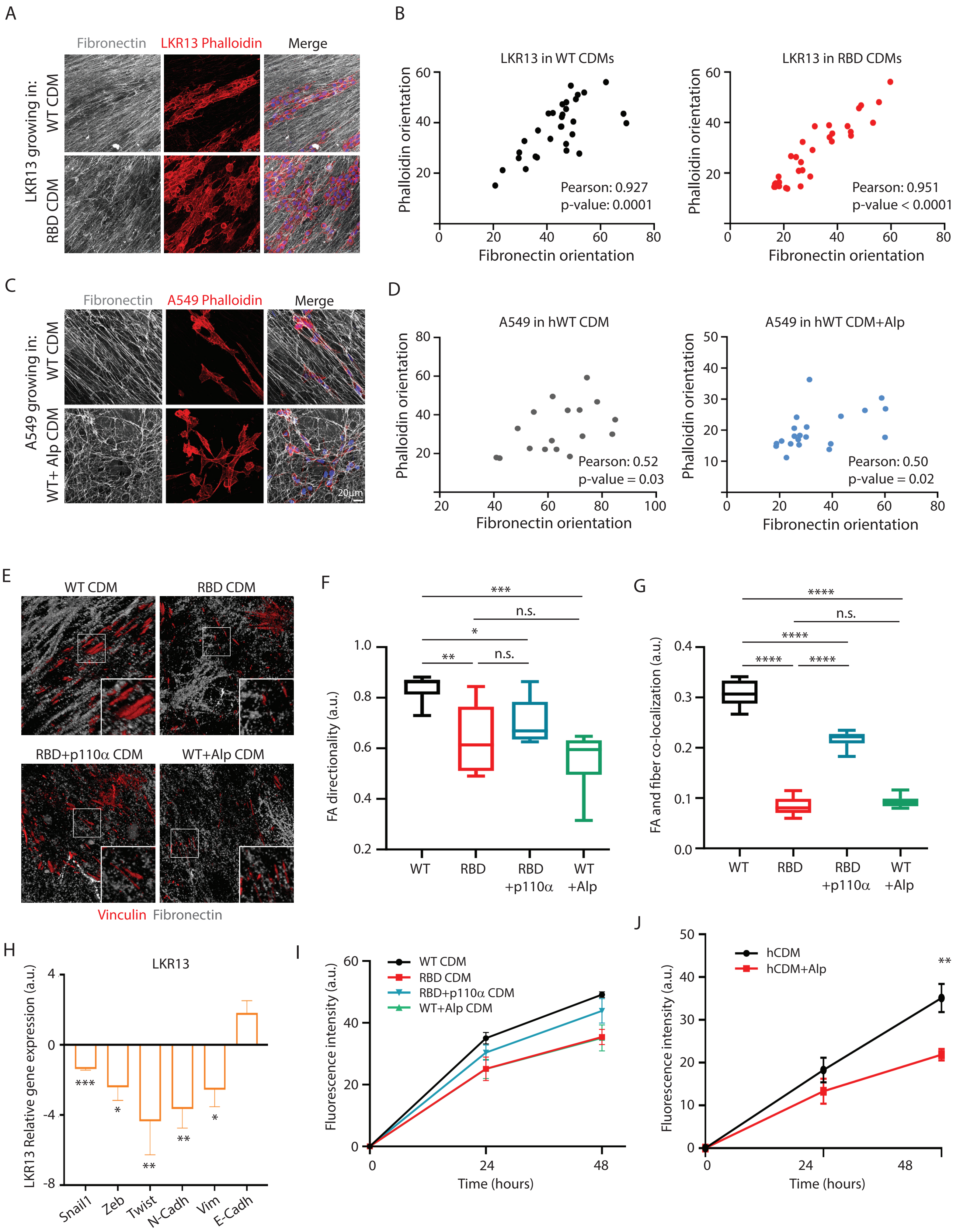

### Supplementary Figure 7S

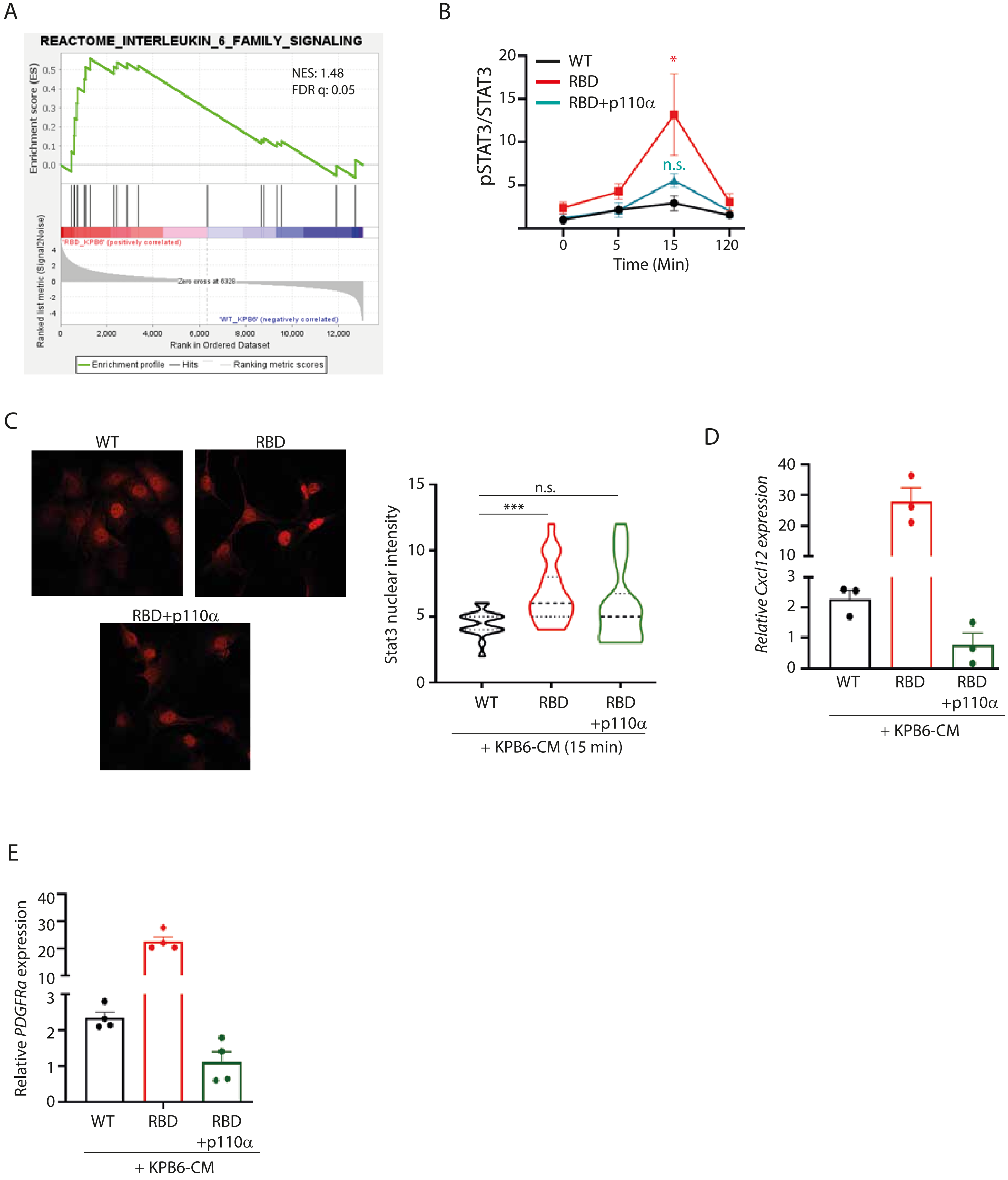
